# The trypanosome mRNA decapping enzyme ALPH1 prefers caps without m^7^G methylation and produces diphosphate-RNA

**DOI:** 10.64898/2026.01.29.702356

**Authors:** Leticia Pereira, Dawid Arkadiusz Dzadz, Paula Andrea Castañeda Londoño, Marcin Warminski, Joanna Kowalska, Nicole Seifert, Silke Braune, Fabiola Holetz, Martin Zoltner, Susanne Kramer, Maria Wiktoria Górna

## Abstract

5′ends of eukaryotic mRNAs are protected by the m^7^G cap, connected to the mRNA via a three-phosphate-bridge. In mRNA decay, the pyrophosphate bond between the α and β phosphate is cleaved by the nudix hydrolase DCP2. Uniquely among eukaryotes, Kinetoplastida lack DCP2 and instead employ the ApaH-like phosphatase ALPH1 for mRNA decapping. ALPH1 consists of an unstructured N-terminus, a catalytic domain and a structured C-terminus that mediates ALPH1 dimerisation. Here, we have analysed *Trypanosoma brucei* ALPH1 in greater detail. We find that the enzyme has broad substrate specificity and accepts different cap types and even cap analogues. Strikingly, cap-analogues and RNAs without the m^7^G-methyl group are turned over significantly faster than m^7^G methylated substrates. Moreover, all methylated and non-methylated cap analogues tested, with at least one additional nucleotide 3′ to the NpppN moiety are cleaved at the β-γ pyrophosphate bond, producing the equivalent to a 5’ diphosphate-RNA. While the presence of the ALPH1 C-terminal domain is essential for cell viability and increases enzyme activity in vitro, substrate preferences are determined solely by the catalytic domain. Altogether, these ALPH1 enzymatic properties exhibit intriguing differences to the canonical eukaryotic decapping enzyme DCP2, which we critically discuss and which potentially have biotechnological applications.

## INTRODUCTION

Eukaryotic mRNAs are protected from degradation by the presence of a 5′cap, an 7-methyl guanosine (m^7^G) that is 5′-5′attached to the first transcribed nucleotide via a bridge of three phosphates (1, 2). Dependent on the organism, further methylations can be present on the ribose or base moieties of the first few nucleotides. mRNA decay usually starts with the removal of the poly(A) tail by deadenylases and can then proceed either 3′to 5′ (exosomal decay) or 5′to 3′. In 5′-3′decay, the cap is removed by the nudix hydrolase DCP2, in a complex with DCP1 and further proteins, most notably Pat1, Lsm1-7 and the enhancers of decapping EDC1-3, and, in metazoans, EDC4. The cleavage occurs at the α-β pyrophosphate bound and produces a 5′ monophosphorylated RNA, that is then further degraded by the exoribonuclease XRN1.

While DCP2 and most of its associated proteins are highly conserved in eukaryotes, they are markedly absent in Kinetoplastida, a group of eukaryotes that includes important human pathogens causing Leishmaniasis, Chagas disease and African Trypanosomiasis. Instead, this group of flagellated protozoans encode the ApaH like phosphatase ALPH1, a bacterial-derived pyrophosphatase of the phosphoprotein phosphatase (PPP) family. In *Trypanosoma brucei*, ALPH1 is the functional equivalent to DCP2 (3), and it is safe to assume, that this function is conserved across the Kinetoplastida (4). The recruitment of an ApaH-like phosphatase for mRNA decapping in Kinetoplastida correlates with a rather specialised way of mRNA processing in this group of protists: mRNAs are transcribed as long polycistronic precursors spanning ten to hundreds of genes, subsequently trans-spliced to the capped, 39 nucleotide long miniexon (ME) sequence of the spliced leader RNA (SL RNA), in a process that is coupled to the adenylation of the upstream gene (5–8). Moreover, the 5′cap provided by the SL RNA is of the rather unusual, heavily methylated type 4: the first four transcribed nucleotides (AACU) carry ribose 2’-O methylations and there are base methylations on the first (m_2_^6,6^A) and fourth (m^3^U) position (9, 10). The hallmark of eukaryotic caps, the m^7^G-methylgroup at the guanosine upstream of the three-phosphate bridge, is conserved in trypanosomes. Thus, a common scheme of all trypanosome mRNAs is the type4 capped 39 nucleotide long miniexon sequence at their 5′ends.

*T. brucei* ALPH1 is a 79 kDa protein and consists of an N-terminal domain, a catalytic domain and a C-terminal domain, that are roughly equal in size (3). The N-terminal domain, predicted to be unstructured, mediates the localisation of ALPH1 to the posterior pole of the cell (11), and is rather poorly conserved across the Kinetoplastida and even absent in ALPH1 of *Trypanosoma grayi* and *Leptomonas pyrrhocoris* (4, 11). Moreover, both procyclic and bloodstream forms can persist with an N-terminally truncated ALPH1 with only a slightly reduced growth rate, indicating that neither the N-terminus, nor posterior pole localisation of ALPH1 is essential for ALPH1 function, at least not in cultured cells (11). The C-terminus, in contrast, is more conserved, required for ALPH1 dimerisation as well as for interactions with proteins forming the ALPH1 decapping complex (11). A range of ALPH1-interactors have been identified by proximity labelling and co-immunoprecipitation of wild type ALPH1 and truncated versions lacking either the C-terminal domain, the N-terminal domain, or both (11). Five of these proteins share the posterior pole localisation with ALPH1, including the *T. brucei* Xrn1 homologue XrnA and a CMGC-like kinase, that both require the ALPH1 C-terminus for their interactions with ALPH1 (3, 11).

The usage of an ApaH like phosphatase in mRNA decapping is unique and the enzymatic properties of *T. brucei* ALPH1, in particular substrate range and preference, remain poorly characterised. Moreover, initial *in vitro* decapping assays indicated, that the product of ALPH1 is not a 5’ monophosphorylated RNA, because it is resistant to Xrn1 exoribonuclease activity (3), indicating that ALPH1 cleaves the pyrophosphate bond at a non-conventional position. Here, we set out to investigate ALPH1 activity *in vitro* by decapping assays, using a wide range of substrates and ALPH1 truncations. ALPH1 cleaved all mRNA substrates and cap analogues tested, but, unexpectedly, displayed clear preference for non-physiological cap types lacking m^7^G methylation. Moreover, in all cap analogue variants that we tested, ALPH1 produced the (equivalent to) a 5’ diphosphorylated RNA, cleaving the pyrophosphate bond between the β and γ phosphate, with m^7^GpppG being the only exception. While absence of the C-terminal domain massively decreases ALPH1 activity *in vitro* and prevents cell growth *in vivo, it* does not affect ALPH1 substrate preferences.

## RESULTS

### Overview of mRNA decapping assays used in this work

We set out to biochemically characterise the enzymatic properties of *T. brucei* ALPH1 in greater detail using *in vitro* mRNA decapping assays. Dependent on the respective substrate and specific question, we employed different methods to detect and accurately quantify the products of ALPH1 activity. These are summarized below, listing their advantages, disadvantages, and controls (Figure 1).

**Figure 1:**
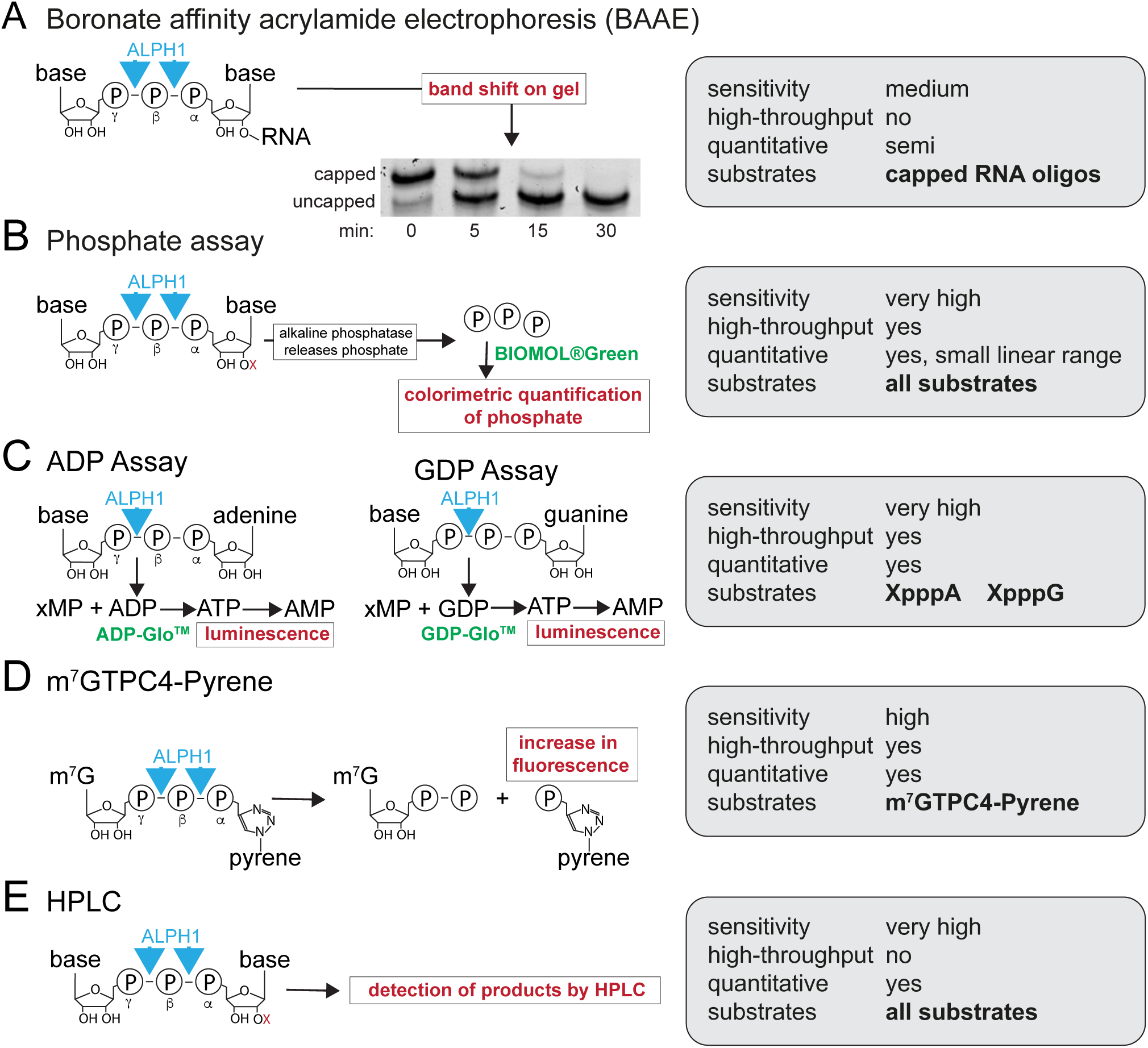
Overview of all mRNA decapping assays used in this work. Five different assays were used to detect the products of ALPH1 decapping activity. All are summarised schematically, highlighting the different read-outs (red text colour) and assay trade names (green text colour). The basic parameters of each assay are summarised in a grey box on the right.

In boronate affinity acrylamide electrophoresis (BAAE) (Figure 1A), capped and uncapped RNA oligos are distinguished by a band shift on a boronate-acrylamide gel (12). This assay was previously used to detect TbALPH1 activity (4) and incubation with an inactive mutant of ALPH1 (D278:N) causes no bandshift, ruling out unspecific cleavage (Figure S1A in supplementary Figures). The BAAE assay works with RNA and is independent of additional enzymes. However, it is low throughput, lacks sensitivity and thus provides only semi-quantitative data. An additional problem further limiting quantitative conclusions is that commercially-derived RNA oligos contain a certain fraction of uncapped oligo, that varies between batches and cap types.

The phosphate assay (Figure 1B) colourimetrically measures the accessible, terminal phosphate that is liberated by alkaline phosphatase from the ALPH1 cleavage products. The assay is high throughput, highly sensitive and has the major advantage, of being substrate-independent, thus, applicable to all types of cap analogues and capped RNA. However, it is limited by a narrow linear substate range of only ∼20-fold (0.4 µM to 8 µM substrate, equalling 1.2 µM to 24 µM phosphate).

In the ADP or GDP coupled enzymatic assays (Figure 1C), ALPH1 produces ADP or GDP from dinucleotide cap analogues, which are in a second step enzymatically transformed into ATP, fuelling luciferase bioluminescence as read-out. This assay is highly sensitive with a wide linear range (15 µM to >500 µM ADP, 0.5 µM to 25 µM for GDP), but restricted to quantification of ADP and GDP from dinucleotide-substrates. Importantly, ADP itself is not a substrate for ALPH1 (Figure S1B in supplementary material).

In the m^7^GTPC_4_-Pyrene assay (Figure 1D), the fluorescent turn-on probe m^7^GTPC_4_-Pyrene (m^7^G-Py) is used as a substrate. Cleavage triggers an increase in fluorescence of the cap-adjacent pyrene moiety used as a read-out for decapping activity (13, 14). It remains unclear to what extent the cleavage site position (β-γ versus α-β) would affect the increase in Py fluorescence, leaving us to infer a cleavage pattern as observed for m^7^G cap analogues (see below). However, we detect the expected ALPH1 dependent fluorescence increase, and the phosphate assay shows that m^7^G-Py is cleaved by ALPH1ΔN with medium efficiency in comparison to other cap analogues (Figure S1C in supplementary material). The m^7^GTPC_4_-Pyrene assay is high-throughput and, importantly, independent of coupled enzymes, which makes it our method of choice for inhibitor studies.

Lasty, reversed-phase high-performance liquid chromatography (RP HPLC) was used to directly detect the decapping products separated by their hydrophobicity and length. In HPLC, most ALPH1 products are detectable in a quantitative way, with the exception of large RNA moieties.

We used recombinant ALPH1ΔN for most experiments, as wild type ALPH1 is more difficult to produce and loses activity quickly even when stored at −80°C. ALPH1ΔN/- trypanosomes are viable (11), indicating that the loss of the N-terminus has no major effect on enzyme characteristics.

### Effects of temperature, pH and ions on ALPH1 mRNA decapping activity

To define the basic enzymatic parameters of ALPH1, we caried out BAAE assays with ALPH1ΔN, varying temperatures, pH values and metal ions. In all assays, we used the 39 nucleotide cap0 RNA oligo with the miniexon sequence as a substrate (4, 11) and the previously described decapping buffer (100 mM NaCl, 50 mM Tris-HCl, 10 mM MgCl_2_, 1 mM DTT, pH 7.9), with respective modifications for the pH and ion experiments.

The decapping rate was highest at 37°C and decreased at higher and lower temperatures; there was still robust activity at 27°C, the temperature optimum cultivation for procyclic trypanosomes (Figure 2A). ALPH1ΔN was active over a wide pH-range (2–10), with an optimum at pH 8 (Figure 2B). No decapping activity was seen in the absence of the enzyme at either pH, excluding the possibility of auto-hydrolysis driven by extreme pH values (Figure S2A in supplementary material).

**Figure 2:**
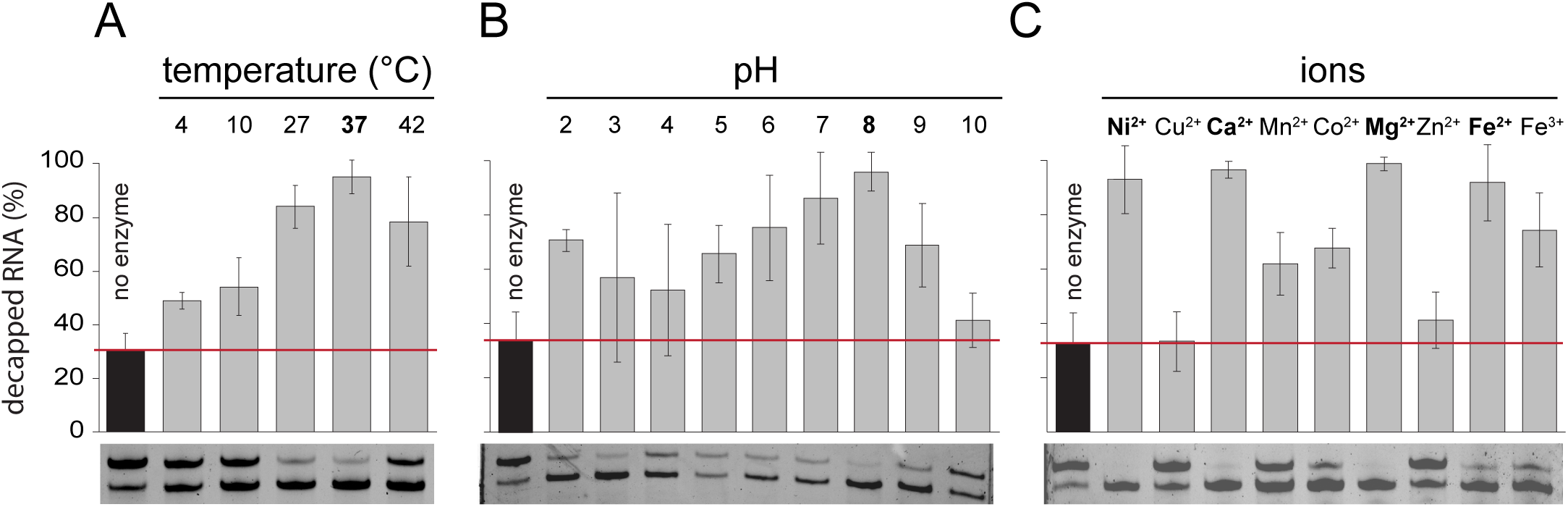
Effect of temperature, pH and divalent cations on ALPH1 activity. BAAE assays were performed with 25 nM ALPH1ΔN and 390 nM RNA oligo (cap0, 39 nucleotide miniexon sequence) in decapping buffer at different **(A)** temperatures, **(B)** pH values, **(C)** or in the presence of different ions. All reactions were performed for 1 hour at 37°C (or the respective temperatures) in decapping buffer (modified to the respective pH or, with MgCl_2_ replaced by 1 mM of the indicated divalent metal cations, NiSO_4_, CuCl_2_, CaCl_2_, MnCl_2_, CoCl_2_, MgCl_2_, ZnCl_2_, FeSO_4_ or FeCl_3_). All assays were performed in triplicate and the percentage of decapped RNA oligo is indicated as average with the error bar representing the standard deviation. An assay omitting the enzyme served as control, and the red line marks the background-level of decapped RNA, already present in the RNA oligo preparation. One representative gel is shown for each assay.

When testing metal ion-dependency, we were surprised to discover that ALPH1ΔN was active in decapping buffer without MgCl_2_, thus, without any divalent ion (Figure S2B in supplementary material). It is unlikely that ALPH1ΔN functions without a divalent ion cofactor, as it is a conserved metallo-phosphatase of the phosphoprotein phosphatase (PPP) group and the mutation of D278, which is involved in chelating the metal ion in the catalytic center, fully abolished enzyme activity (Figure S1A in supplementary material). Instead, the supportive ion likely copurified with the enzyme and bound with high affinity: many metalloenzymes require a metal-ion cofactor for efficient and correct folding (15), including ApaH of *S. flexneri* (16). When testing different ions at 1 mM concentrations, we found high ALPH1 activity with Mg^2+^, Ca^2+^, Ni^2+^, Fe^3+^ Fe^2+^, and to a slightly lesser extent with Mn^2+^ and Co^2+^, indicating that these ions either support ALPH1 decapping activity or fail to replace the ion that is already bound (Figure 2C). Cu^2+^and Zn^2+^, had inhibitory effects; these ions are known to bind stronger to proteins than Mg^2+^ and may out-compete Mg^2+^ when given at such unphysiologically high concentrations (17). In conclusion, ALPH1 is active over a wide temperature and pH range and is supported (or at least not substantially inhibited) by Mg^2+^, and its ion cofactor appears to be tightly bound.

### ALPH1 displays low sensitivity towards phosphatase inhibitors and EDTA

Next, we tested the sensitivity of ALPH1 to known inhibitors. ALPH1 belongs to the PPP group of phosphatases and we therefore tested two classical phosphatase inhibitors, sodium orthovanadate and okadaic acid, as well as EDTA, which inhibits phosphatases by chelating the metal ion cofactors.

As a positive control, we first tested the phosphatase activity of commercially available protein phosphatase 1 (PP1) on *p*-nitrophenyl phosphate (pNPP) using colorimetry to quantify the cleavage product *p*-nitrophenol (Figure 3A). PP1 was readily inhibited by all three compounds, with IC_50_ values of 0.02 mM for sodium orthovanadate (Na_2_VO_3_), 2.3 µM for okadaic acid and 2.1 mM for EDTA. For ALPH1ΔN, we used m^7^GTPC_4_-Pyrene as a direct substrate, to avoid any unspecific effects on enzymes unrelated to ALPH1 (Figure 1D). We found no inhibition of ALPH1ΔN by up to 40 µM okadaic acid (Figure 3B). Both sodium-orthovanadate and EDTA had an inhibitory effect on ALPH1ΔN, but with very high IC_50_ values of 0.23 mM and 17 mM, respectively (Figure 3B). Thus, ALPH1ΔN responds poorly to classical phosphatase inhibitors. The insensitivity towards EDTA supports our above-described model of a tightly-bound divalent ion in the catalytic centre of ALPH1, protected from chelation by EDTA.

**Figure 3:**
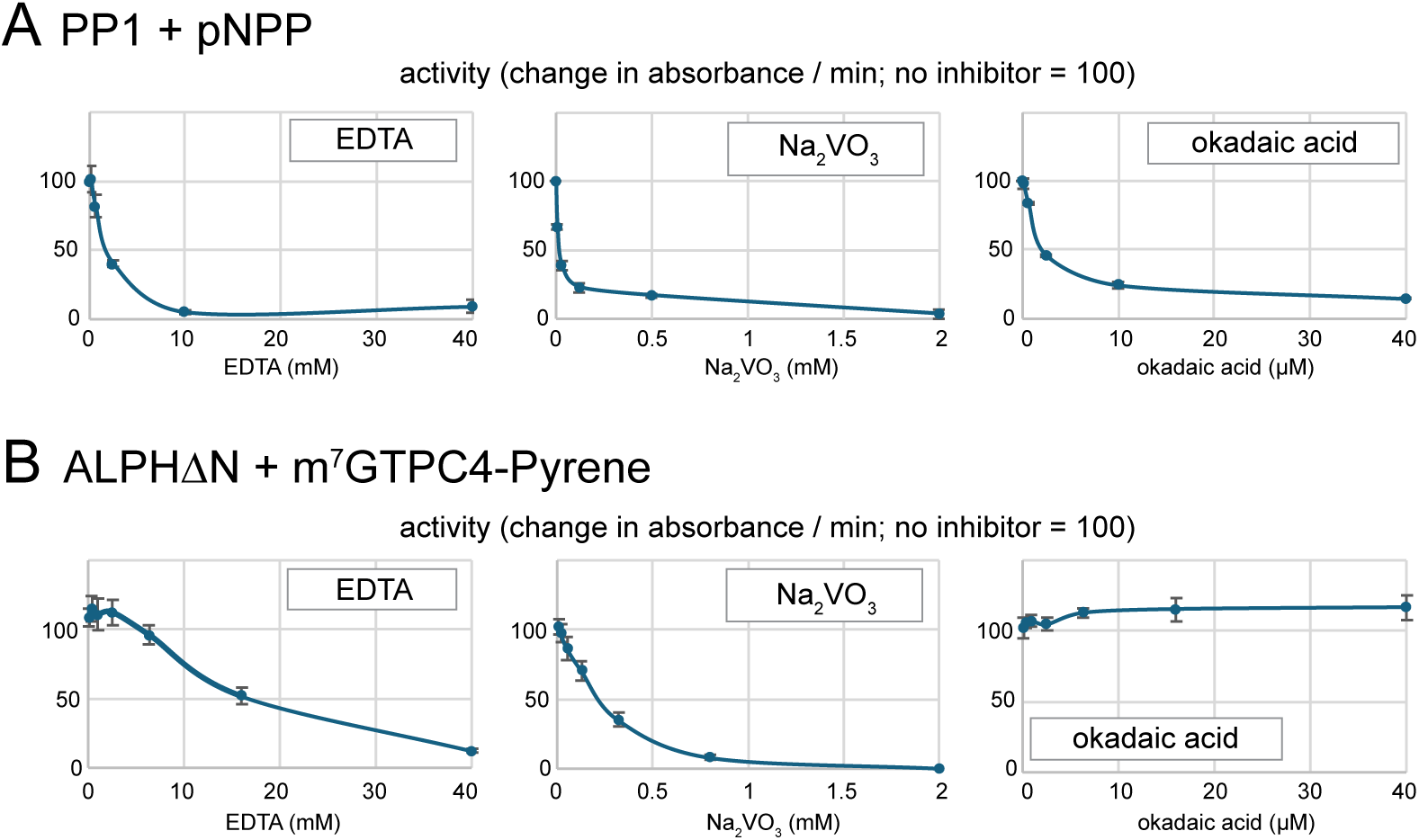
Phosphatase inhibitor profile. **A)** Phosphatase assay with Protein Phosphatase-1 catalytic subunit (PP1). 10 mM pNPP was used as substrate and the production rate of the product, para-nitrophenol, was quantified colorimetrically. The assay was performed in the presence of a titration of the phosphatase inhibitors sodium vanadate, okadaic acid or the chelator EDTA and 0.02 U/µl of PP1. All experiments were in triplicates and average values are presented, with error bars representing standard deviations. **B)** ALPH1ΔN activity was measured using the m^7^GTPC4-Pyrene decapping assay, in the presence of the same titration of phosphatase inhibitors as in A and 500 nM of ALPH1ΔN. The production rate of the product was calculated, and averages of triplicate experiment are presented, with error bars representing standard deviations.

The low sensitivity of ALPH1 towards phosphatase inhibitors correlates with a poor activity of ALPH1 towards pNPP, the classical substrate of *in vitro* phosphatase assays. At 0.1 µM ALPH1ΔN, a concentration sufficient for complete turnover of cap-analogs (at the same substrate concentration and time period), there is almost no pNPP product detectable, and even at 0.8 µM, pNPP turnover is poor (Figure S3 in supplementary material).

In summary, even though ALPH1 belongs to the PP1 family, it does not share substrate preferences or inhibitor sensitivity with protein phosphatases.

### ALPH1 cleavage is not specific to the type 4 cap or the miniexon sequence

The BAAE assays above used a cap0 RNA oligo as a substrate. Trypanosome mRNAs have a cap4, and we therefore tested, whether this would be a better substrate for ALPH1. A cap4-RNA is not commercially available, and we therefore used a slightly modified cap4 (cap4*), lacking the methyl group at the uracil, which is the 4^th^ base after the cap (Figure 4A). Note that this oligo preparation was not clean and contained approximately 50% uncapped RNA. We compared mRNA decapping activity of ALPH1ΔN towards RNA oligos capped with a cap0 or a cap4* and found that ALPH1ΔN was active towards both oligos (Figure 4B). We next varied the sequence of the cap0 oligo, using a shuffled sequence of the miniexon. In a second experiment, we used a cap0 oligo modified to have G instead of A as the first “transcribed” base, followed by the miniexon sequence. Neither the change in RNA sequence, nor the change to a m^7^GpppG cap caused any substantial change in mRNA decapping activity (Figure 4C). Our data do not support any preference of *T. brucei* ALPH1 towards the trypanosome-specific cap4 type.

**Figure 4:**
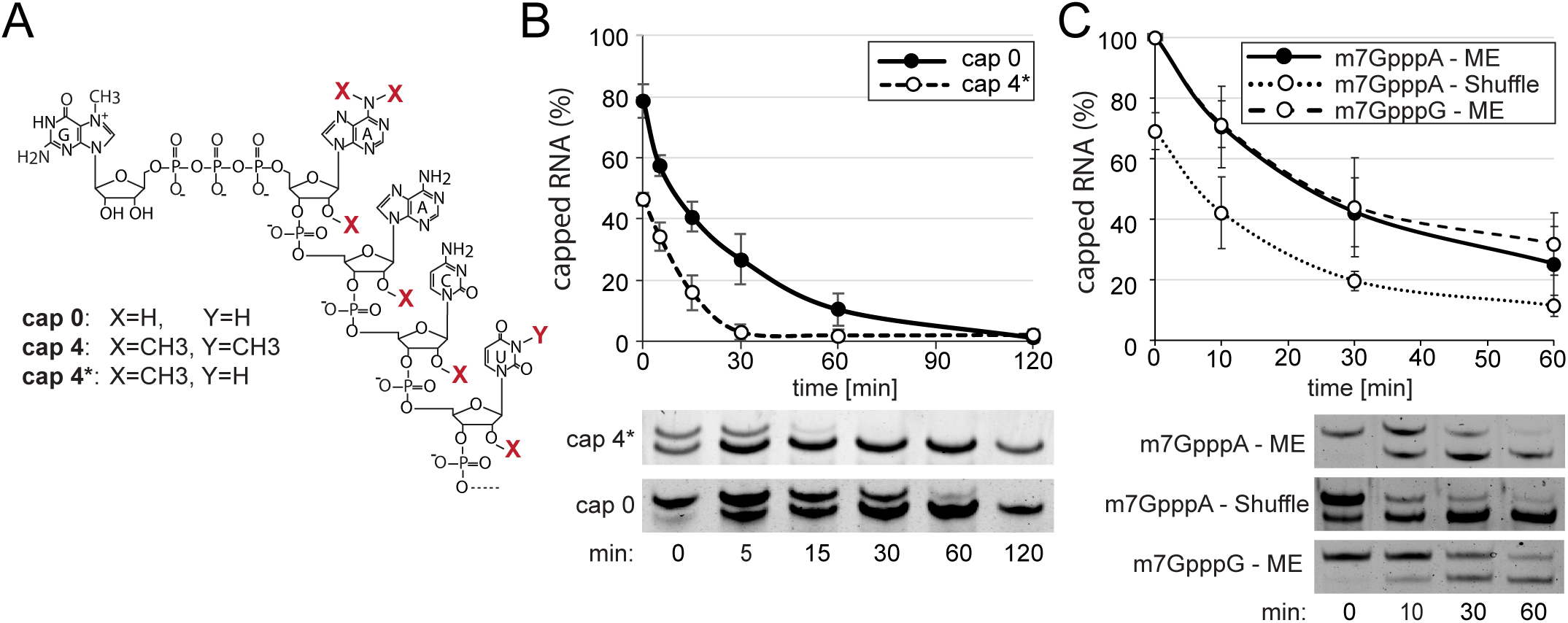
Different cap types and RNA sequences. BAAE-assays were performed with ALPH1ΔN and a range of different RNA substrates. The percentage of capped RNA was quantified from three independent experiments. Average values are shown, with error bars representing standard deviations. One representative gel is shown for each experiment. **A)** Chemical structure of cap0, cap4 and cap4* scaffolds. **B)** Dependency on cap type: A decapping assay with ALPH1ΔN was performed over a time-course of 120 min, using either cap0 or cap4* as a substrate. **C)** Dependency on sequence: A cap0 oligo with a shuffled miniexon sequence (m^7^GpppA-GUUUUAUCAGAAUAGAAUCUAGACCUUUAUGUUUAACC) and a cap0 oligo with an m^7^GpppG cap instead of m^7^GpppA were used as substrates for decapping assays with ALPH1ΔN. The standard cap0-ME oligo served as a control.

### ALPH1 prefers substrates without the m^7^G -methyl group

Every eukaryotic mRNA, including trypanosome mRNAs, has a methyl-group at nitrogen position 7 of the guanosine (m^7^G cap). When we compared decapping of a 150 nucleotide long RNA capped with either m^7^G and just G, we found, surprisingly, that ALPH1ΔN prefers the RNA with the non-methylated cap over its physiological substrate (Figure 5A).

**Figure 5:**
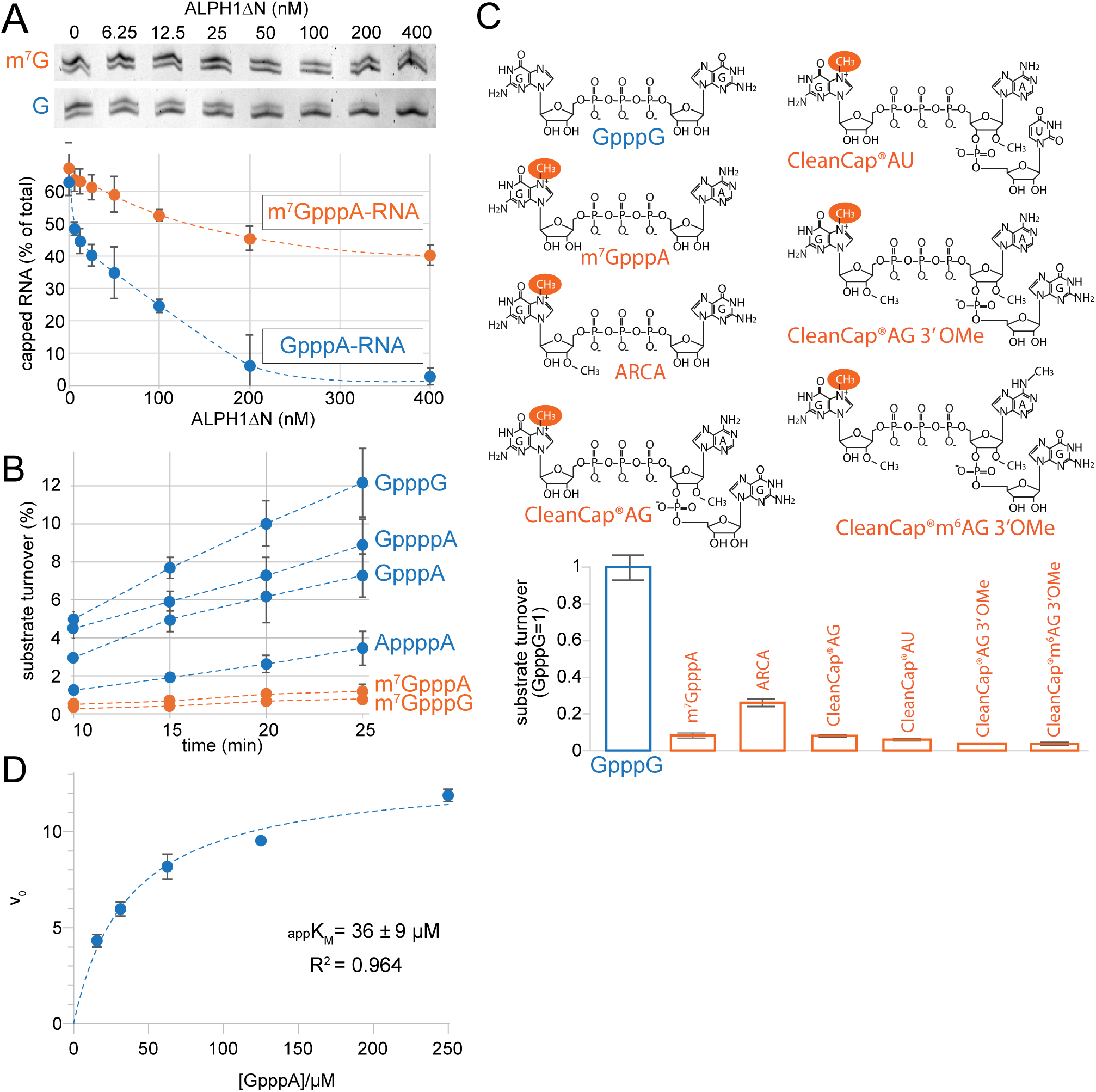
ALPH1 substrate preference. **A)** A BAAE-assay was performed for 30 min, with ALPH1ΔN ranging from 0 to 400 nM and 51.2 nM of a 150 nt long RNA substrate with a type 0 cap, either with the m^7^G methyl-group or without. The assay was done in triplicates, using substrate of two different preparations. The % of capped RNA was quantified and is presented, with error bars presenting standard deviations. Representative gels of one experiment are shown. **B)** A phosphate-assay was performed with ALPH1ΔN (1.5 nM) and 50 µM of different cap analogues, over a 25 min time-course. Data are presented as average substrate turnover (in %) from three independent experiments with the standard deviation indicated by error bars. **C)** A phosphate-assay was performed with ALPH1ΔN (1.5 nM) and 50 µM of different cap analogues, for 25 minutes. The substrate turnover was measured and is shown in relation to the phosphate-release of the best ALPH1 substrate, GpppG. Importantly, the substrate turnover was always much less than 20%, except for GpppG, where 20% of the substrate were consumed. **D)** Determination of the apparent K_M_ (appK_M_) for ALPH1ΔN (21 nM) for the GpppA substrate via phosphate assay. The multiple-turnover reaction was over 15 min and appK_M_ and R^2^ values are indicated below the curves. For a replicate analysis see Figure S6.

Therefore, to analyse the substrate preferences of ALPH1 in greater details and in a more quantitative way, we switched to the phosphate assay (Figure 1B) and used cap-analogues as substrates. We included cap-analogues with and without the *N7*-methyl group at the guanosine, namely: GpppA, GpppG, m7GpppA, m7GpppG, AppppA and GppppA, using substrate-excess conditions with substrate turnover of <15% at the last timepoint (Figure 5B). All cap analogues were accepted as ALPH1ΔN substrates, but the turnover rates differed. The three best substrates were the non-methylated, guanosine-containing cap-analogues GpppG, GppppA and GpppA. Slightly less well-accepted, but still better than the methylated cap-analogues, was AppppA, indicating that ALPH1 activity is not restricted to substrates with three phosphate bonds. Of all these substrates, the cap-analogues with m^7^G methylations had the lowest turnover rates. We tested five further cap analogues with m^7^G-methylations. These included the anti-reverse cap analogue (ARCA, m_2_ ^7,3’-O^GP G) as well as four cap analogues of the CleanCap® series that contained an additional nucleotide (Figure 5C). All five cap analogues had low substrate turnover rates, similar to m^7^GpppA and significantly lower than GpppG.

We determined the appK_M_ for ALPH1ΔN with the GpppA substate under multiple-turnover conditions employing the phosphate assay (Figure 5D; Figure S6). The appK_M_ was 36 +/-9 µM, while derived catalytic efficiency values were high (appk_cat_ = 632 min^-1^). This suggests that the substrate binding step (and potentially the product release) is dominating turnover. We thus expressed parameters as apparent values as multiple-turnover kinetics may not ideally reflect this scenario.

In conclusion, ALPH1 prefers non-methylated cap substrates, which is unexpected and may reflect the evolutionary origin of this enzyme from the bacterial ApaH, which prefers (unmethylated) diadenosine tetraphosphate. Our kinetics analyses suggest a low substrate affinity, which under physiological conditions could be enhanced by interaction with other components of the decapping complex, analogous to the action of enhancers of decapping found in the DCP2 complex.

### ALPH1 cleavages leaves a diphosphorylated product

One only partially answered question was which of the two pyrophosphate bonds is preferentially cleaved by ALPH1. We had previously shown XRN1 exoribonuclease activity resistance of ALPH1-cleaved SL-RNA, which could be resolved by the addition of RNA-5′polyphosphatase, indicating that the decapped mRNA, unusually, does bear more than a monophosphate (3).

We first determined the cleavage site positions of the cap analogues GpppA and m^7^GpppA. We used the phosphate assay, to establish conditions that caused the decapping reaction to go to completion (Figure 6A, left). With these conditions, we performed an ADP assay. This assay should produce 100% ADP, when the pyrophosphate bound next to the guanine is cleaved, and respectively less, if the other cleavage site is used. We observed almost 100% ADP production with GpppA, but only about 50% with m^7^GpppA, indicating that the presence of the m^7^G methyl group shifts the cleavage site preference (Figure 6A, right). Next, we compared GpppG and m^7^GpppG in the same way. m^7^GpppG is a poor substrate for ALPH (compare Figure 5B) and we failed to get the reaction to completion, even with half the substrate concentrations (Figure 6B, left). Still, 70% of m^7^GpppG was cleaved, but almost no GDP was produced, indicating that m^7^GpppG is preferentially cleaved at the pyrophosphate bond between the α and β phosphate (Figure 6B, right). Thus, for dinucleotide cap analogues, the data are consistent with a preference of ALPH1 to cleave adjacent to a guanine rather than adenosine and distant to the m^7^G-methyl group. For m^7^GpppG, both preferences add up to the same cleavage site, while for m^7^GpppA these preferences are conflicting, offering a possible explanation for the differences in cleavage site preference between the two substrates.

**Figure 6:**
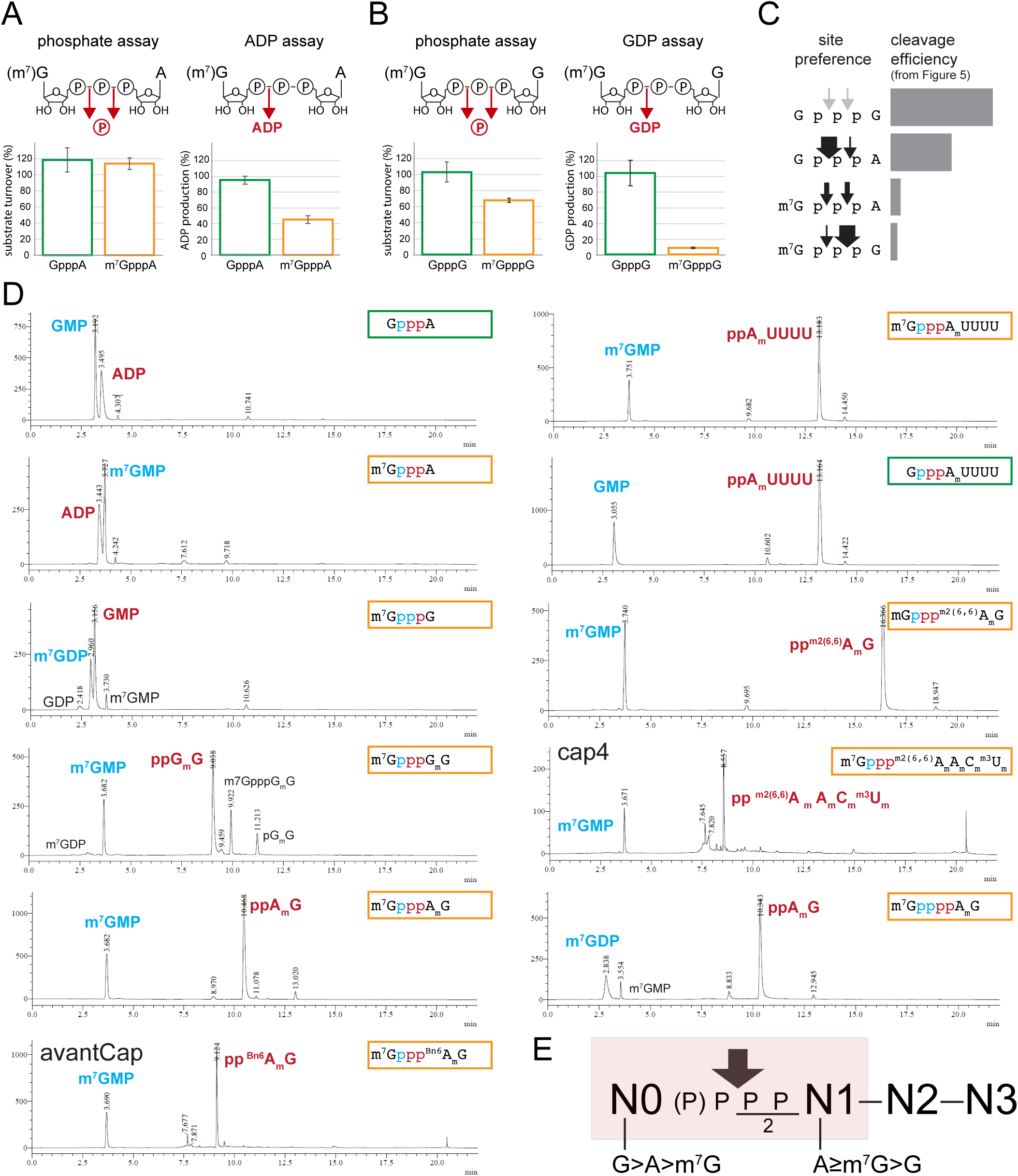
ALPH1 cleavage site preferences. **A)** Using the phosphate assay, decapping reactions were established for ALPH1ΔN that resulted in the reaction being complete for both, the m^7^GpppA and GpppA substrate. These conditions were 15 µM substrate, 0.11 µM ALPH1 in standard decapping buffer for 1 hour. Under these conditions, ADP release was quantified from both, m^7^GpppA and GpppA, with the ADP assay. Data of three independent experiments are shown, with the standard deviation indicated by error bars. **B)** The experiment was repeated for m^7^GpppG and GpppG, quantifying GDP using the GDP assay. Note that even though the substrate concentration was halved, full completion of the reaction could not be achieved for m^7^GpppG. **C)** Summary of relative cleavage site preferences, as determined by the ADP and GDP assay **D)** The HPLC assay was used to determine the cleavage products for a range of cap analogues. The decapping assay was done with 50 µM cap analogue and 1 µM ALPH1ΔN for 1 hour at 25°C. The HPLC profiles are shown, and the main products are labelled. The substrate is indicated in the frame, and the observed cleavage site position indicated by the change in text-colour of the phosphate group. **E)** Summary of cleavage site preference and preferred nucleosides of ALPH1.

Next, we used HPLC to detect the cleavage products from the same cap analogues (Figure 6D). For m^7^GpppG and GpppA, the cleavage site preferences were identical to the ones determined by the ADP/GDP assays: both substrates were cleaved next to the non-methylated G, resulting in the equivalent of a monophosphate-RNA for m^7^GpppG and of diphosphate-RNA for GpppA. For m^7^GpppA, the HPLC showed preferential cleavage into ADP and m^7^GMP rather than the equal use of both cleavage sites as observed in the ADP assay. This small discrepancy between the data may be caused by differences in the assay conditions. We used HPLC to detect the cleavage products of eight additional cap-analogues that all had additional nucleotides and methylations, namely m^7^GpppG_m_G, m^7^GpppA_m_G, m^7^Gppp^m2^(^6,6^)A_m_G (18), m^7^GppppA_m_G (19), the trypanosome cap4 m^7^Gppp^m2^(^6,6^)A_m_A_m_C_m_^m3^U_m_ (20), m^7^Gppp^Bn6^A_m_G (21) and m^7^GpppA_m_UUUU and GpppA_m_UUUU. All these cap analogues were cleaved by ALPH1ΔN into products that corresponded to a diphosphate-RNA and not to a monophosphate-RNA. Importantly, this was independent of the methylation status and included artificial cap analogues like the AvantCap m^7^Gppp^Bn6^A_m_G (21). Thus, the presence of at least one additional nucleotide is sufficient to shift the cleavage site preference of ALPH1ΔN towards the β-γ pyrophosphate bond. For example, while m^7^GpppG is cleaved into m^7^GMP and GDP, m^7^GpppG_m_G is cleaved mainly to m^7^GDP and pG_m_G. Interestingly, even the cap analogue with a 4-phosphate-bridge, m^7^GppppA_m_G, is cleaved into the equivalent of a diphosphate-RNA. This suggests that the phosphate-groups are “counted” from the RNA-moiety rather than from the cap nucleotide at the 5′end.

In conclusion, we find that cap-analogues are cleaved into the equivalent of a diphosphate-RNA, and this seems independent of the methylation status of riboses and/or bases, type of nucleotide and number of phosphate-groups in the bridge. The only exception is m^7^GpppG, and, possibly partially m^7^GpppA, but these are unlikely to be physiologically relevant, as the cleavage site changes towards a diphosphate, the moment one or more additional nucleotide is added. A model of the cleavage preference of ALPH1, and its nucleoside-preferences in the respective positions of the cap is shown in Figure 6E and will be discussed below.

### The C-terminal domain of ALPH1 is essential *in vivo* and increases enzyme activity *in vitro* but plays no role in substrate preferences

To this point, all experiments had been performed with the N-terminally truncated version of ALPH1, ALPH1ΔN. This ALPH1 truncation has been confirmed to be functional *in vivo*, (11). In contrast to the N-terminus, the C-terminal extension of ALPH1 is predicted to be structured, is conserved across kinetoplastids, mediates ALPH1 dimerisation and its removal prevents ALPH1 localisation to RNA granules, indicating it may be essential for ALPH1 function (11). We thus tested whether the removal of the C-terminus would still result in a functional enzyme, *in vivo* and *in vitro*, by analysing and comparing a range of ALPH1 truncations (Figure 7A).

**Figure 7:**
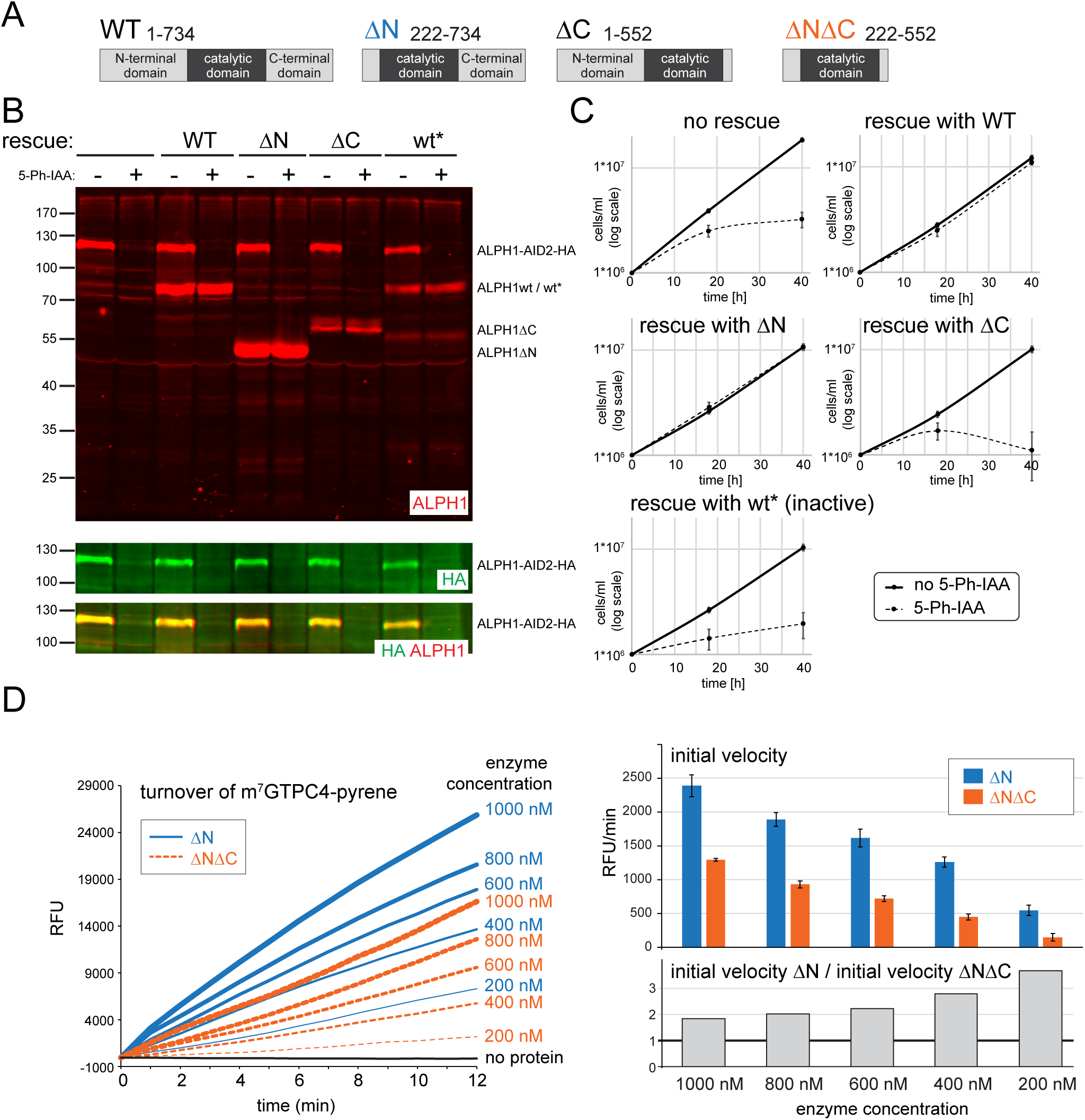
Function of the ALPH1 C-terminus. **A)** To scale schematics of wild type ALPH1 (WT) and the ALPH1 truncation constructs used in this work, ALPH1ΔN, ALPH1ΔC and ALPH1ΔNΔC. **B and C)** ALPH1ΔN, but not ALPH1ΔC can rescue viability of a cell line depleted for ALPH1. One ALPH1 allele was replaced with ALPH1-AID2-3xHA, the other was deleted, in a cell line expressing the necessary components for the degron system (22). Genes encoding wild type ALPH1 and ALPH1 variants were integrated into the tubulin locus for stable expression and the degradation of ALPH1-AID2-3xHA, was induced with 5-Ph-IAA. The successful depletion of ALPH1-AID2-3xHA, and the expression of the ALPH1 variants was controlled by a western blot loaded with protein samples of cells with (2 hours 5-Ph-IAA) and without induction of ALPH1-AID2-3xHA, and probed for ALPH1 and for HA (B). Note that the ALPH1 antiserum was produced against ALPH1ΔN, which is the likely reason for the ALPH1ΔC bands appearing less strong. Growth was monitored with and without induction of the degron; for each cell line, data are averages from three independent clones with standard deviations represented by error bars (C). **D)** The m^7^GTPC4-Pyrene probe was used at 5 µM in decapping assays with ALPH1ΔN and ALPH1ΔNΔC ranging from 200 to 1000 nM. On the left, the substrate turnover is shown over a time-course. On the right, the initial velocities were calculated from a linear regression fitted to the first 6 min of the reaction. RFU=relative fluorescence units. Data are averages of the initial velocities calculated for each repetition, error bars represent standard deviations.

To test the function of the C-terminus *in vivo*, we depleted ALPH1 by the 5-Ph-IAA inducible degron system (22) and monitored, whether a C-terminally truncated ALPH1 variant can rescue the growth phenotype. First, one allele of ALPH1 was deleted and the other was replaced with an ALPH1-AID2 fusion, in the trypanosome cell line that expressed all components for the auxin degron system (22). ALPH1-AID2 protein is degraded within 2 hours of induction with the auxin derivative 5-Ph-IAA (Figure 7B, lane 1 and 2) and this depletion caused the expected growth arrest, within less than 20 hours (Figure 7C, no rescue). Next, wild type ALPH1 and a range of ALPH1 variants were constitutively expressed from the tubulin locus in the same cell line. ALPH1-degron cell lines which expressed wild type ALPH1 or ALPH1ΔN grew normally in the presence of 5-Ph-IAA, while the catalytically inactive ALPH1 D278N mutant (ALPH1 wt*) and ALPHΔC (1–552) were unable to rescue the growth phenotype (Figure 7B-C). The data confirm that the N-terminus of ALPH1 is not required for growth in culture and show that, in contrast, the C-terminus is essential.

Next, we tested, whether the C-terminus of ALPH1 is necessary for its enzymatic activity. We purified ALPH1ΔNΔC, the isolated catalytic domain. First, we tested ALPH1ΔNΔC in the BAAE assay, using a cap0 and cap4 RNA oligo as a substrates. We found mRNA decapping of both substrates, proving that the C-terminus is not necessary for mRNA decapping per se (Figure S4 in supplementary material). For a more quantitative analysis, we compared the mRNA decapping activities of ALPH1ΔNΔC and ALPH1ΔN using the m^7^GTPC4Pyrene assay. We tested both ALPH1 variants in different concentrations over a time-course and found that, for all concentrations, ALPH1ΔN had a higher substrate turnover rate than ALPH1ΔNΔC (Figure 7D, left panel). The difference in initial between ALPH1ΔNΔC and ALPH1ΔN increased with decreasing enzyme concentrations: at 1000 nM, ALPH1ΔN had a 1.85 fold higher initial velocity than ALPH1ΔNΔC and at 200 nM it was 3.68 fold (Figure 7D, right panel). Given that the C-terminus mediates ALPH1 dimerisation (11), it is possible that ALPH1 dimerisation enhances its activity. The gradual decrease in the differences between ALPH1ΔNΔC and ALPH1ΔN activities with increasing enzyme concentrations may be caused by the more active ALPH1ΔN reaching saturating (substrate-limiting) conditions at a lower enzyme concentration than ALPH1ΔNΔC. Alternatively, ALPH1ΔNΔC may still have some residual (substrate dependent?) ability to dimerise, and such inefficient dimerization would be favoured by high enzyme concentrations. Either way, the data show that the C-terminal domain of ALPH1 is a major contributor to ALPH1 activity.

We next tested, whether the C-terminus is involved in the determination of substrate preferences, using the phosphate assay and the same set of cap-analogues that had been tested in Figure 5. While we needed >10 times more ALPH1ΔNΔC than ALPH1ΔN for comparable cleavage rates, confirming the differences between the two enzyme variants described above, we observed no major differences in substrate preferences. Both enzyme variants preferred caps lacking the methyl group of the m^7^G (Figure S5A-B in supplementary materials). The one minor difference was a switch in the order of the two best substrates, ALPH1ΔN preferred GpppG over GppppA, while ALPH1ΔNΔC had opposite preferences. Next, we tested, whether the C-terminus contributes to the phosphate bond cleavage position. The low activity of ALPH1ΔNΔC prevented a large-scale analysis of cleavage site preferences. Nevertheless, we could establish that ALPH1ΔNΔC cleaves GpppA entirely to ADP, just like ALPH1ΔN (Figure S5C in supplementary materials), indicating that there are no differences in cleavage site preferences between the enzyme variants.

In conclusion, the C-terminal domain of ALPH1 is essential *in vivo* and its presence significantly increases enzymatic activity. However, it does not appear to significantly affect substrate and cleavage-site preferences: these features are solely determined by the catalytic domain.

## DISCUSSION

### Favoring the non-methylated cap - an unexpected preference for an mRNA decapping enzyme

One common feature shared by all eukaryotes is the m^7^G-cap at the 5′end of mRNAs, that protects mRNA from degradation and mediates mRNA specific interactions during processing, translation and decay. Trypanosomes are no exception. However, trypanosomes possess a unique m^7^G cap structure with ribose and base methylations ranging up to the 4^th^ transcribed nucleotide, and employ a unique enzyme for mRNA decapping. It seemed an obvious explanation, that this unusual decapping enzyme had evolved as a special adaptation to the unique cap structure. To our surprise, our *in vitro* data do not support this hypothesis. ALPH1 appears rather poorly adapted to the cap4 but instead prefers substrates that lack even the canonical m^7^G methylation. The trypanosome mRNA decapping enzyme is thus fundamentally different to the canonical decapping enzyme DCP2, which prefers substrates with m^7^G-methylation (25) and an RNA moiety of at least 25 nucleotides (25, 26) and produces monophosphate RNA (27, 28).

### An unusual ALPH1 cleavage site producing diphosphate RNAs

A further particularity of ALPH1 is the unusual cleavage pattern of the pyrophosphate bonds, producing a diphosphate RNA as a product, and not a monophosphate, like DCP2 does. Our previous data had already suggested this, based on the observation that ALPH1 cleavage products were insensitive to Xrn1, but could not formally distinguish between a diphosphate or triphosphate RNA product (3). Our new data indicate that the number of residual phosphate-groups is determined relative to the first nucleotide (N1, Figure 6E) rather than to the cap-nucleotide (N0) as cap analogues with four phosphate-groups give the same diphosphate RNA product. This would indicate that ALPH1 recognises and binds to N1. If we then align all dinucleotide cap-analogues in a way, to define N1 as the nucleoside retaining two phosphate-groups after cleavage, our combined data on cleavage position and cleavage efficiency give a preference for N0 to be a guanosine (G>A>m7G) and for N1 to be an adenosine, or, an m^7^guanosine (A≥m^7^G>G) (Figure 6E). While for dinucleotides, the preferences for N0 and N1 likely determine binding orientation and thus the cleavage site, the presence of a nucleotide at position N2 unequivocally produces (the equivalent of) a diphosphate RNA. This is independent of the type of nucleotide and its modifications and occurs even when the nucleotide at N0 is unfavourable, indicating that anything larger than a dinucleotide can only be oriented in one directed way. In the absence of experimental structural data, only a predicted comparative model of ALPH1 can be used to study the putative RNA-binding interface. A recent study has structurally analysed the binding of Np4-RNA to the bacterial decapping enzyme ApaH and found that ApaH can bind to Np4-RNA in one of two orientations, partially dependent on its nucleoside preferences for the N0 and N1 positions (23). However, the Np4-cap is symmetric and ApaH cleaves exactly in the middle (24), while a triphosphate-bond must be cleaved asymmetrically, and we therefore believe, in consistence with our data, that only a single orientation is used by ALPH1 for longer substrates. ApaH crystal structures reveal electron density for its RNA substrate only at one nucleoside and the four-phosphate bridge (24), suggesting only one predominantly strong nucleoside binding site with a preference for A over G (Figure 8A). Most features of this nucleoside binding site are conserved in ALPH1, such as the aromatic character of Tyr488 (in place of key ApaH residue Trp249, which could be successfully substituted with Phe)and the hydrophobic character of Ala469 (ApaH Ala229), suggesting its full functionality in ALPH1. However, Gly472 might decrease adenine specificity similarly to the effect of the Glu232Ala mutant in ApaH. Based on our observations, it seems likely that dinucleotide analogues are also preferably oriented “downwards” with A bound in the nucleoside binding site of ALPH1, while the two cleavage sites in m^7^GpppA and m^7^GpppG suggest that two orientations are possible – the dinucleotide can be flipped “upwards” i.e. with m*^7^*G outside of the nucleoside binding site, and this is more likely when the other nucleotide is A (Figure 8B). The methyl group introduces a positive charge on N7, which might affect interaction of the base located in the nucleoside binding site with the Pi cloud of Tyr488 – this detail could not be inferred from available structural data as no methylated substrates were used in the ApaH study. However, it has to be noted that even for ApaH the nucleobase preference of the nucleoside binding site was not reflected in the substrate cleavage efficiency. Whenever an additional nucleotide is present, the RNA bound to ALPH1 would always assume an “upwards” orientation with N1 in the nucleoside binding site, and the body of the RNA would follow where the phosphate proceeding N2 can be seen in ApaH model with GppppAU bound (the closest sequence among the determined co-crystal structures to the trypanosome spliced leader). For ALPH1, the “phosphate-counting” implicates definite binding at N1, but is quite likely that additional ALPH1 residues can contact N2, N3 and further nucleotides, contributing to the recognition, binding and positioning of the RNA, possibly also with the help of its essential C-terminal domain that is absent in ApaH. One open question is, how the diphosphate-RNA is further degraded. Our extensive proximity labelling data of ALPH1 and its interactors have not identified another pyrophosphatase. The likeliest scenario is, that XRNA, an integral component of the decapping complex, binding the ALPH1 C-terminus (11), has the ability to degrade diphosphate RNA.

**Figure 8:**
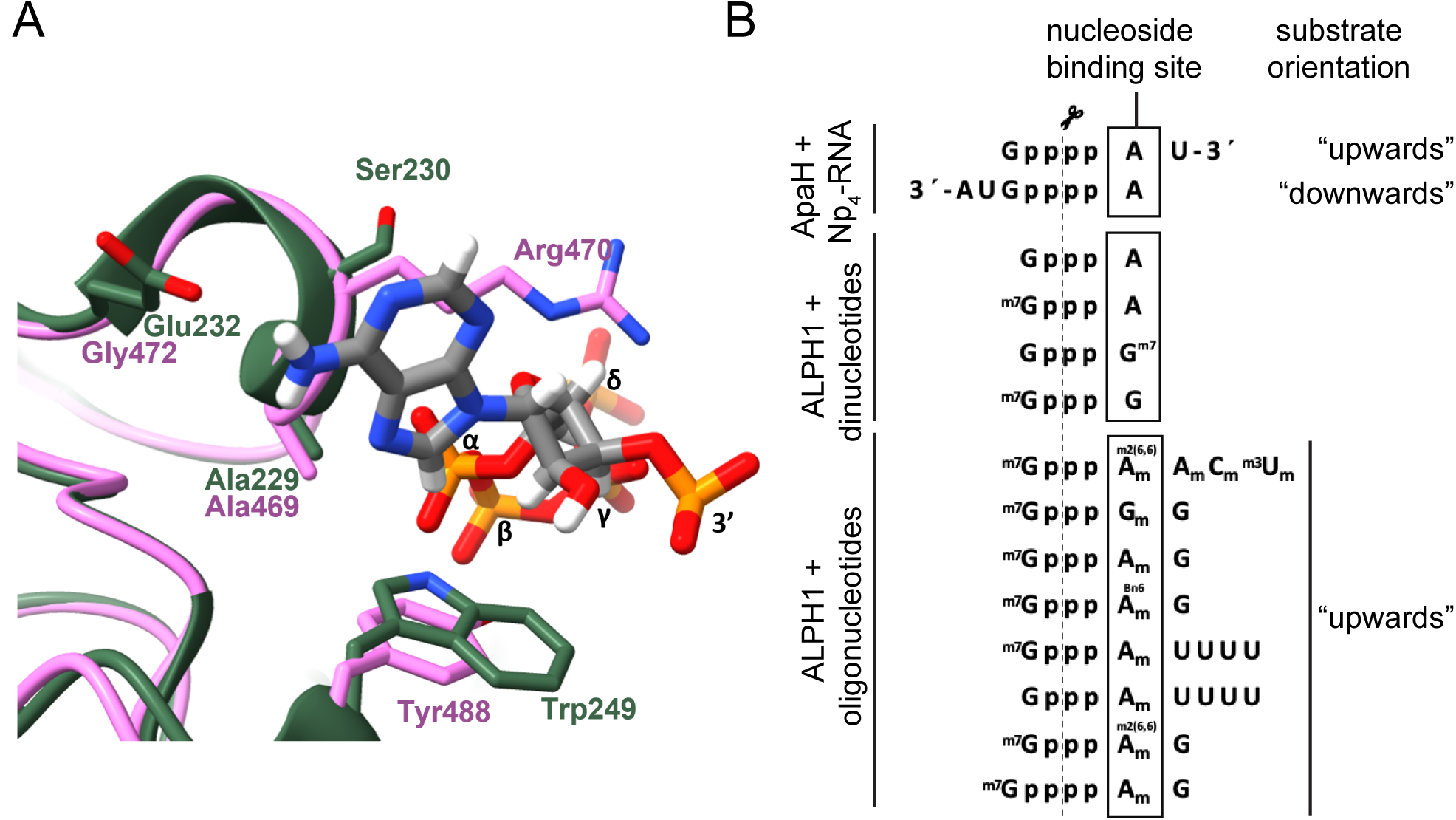
Structural comparison with bacterial ApaH bound to a substrate. **A)** A comparison of a predicted model of ALPH1 (rose) with a crystal structure of *E. coli* ApaH (green, PDB entry 9ooy) bound to GppppAU (gray). Indicated are key amino acids contributing to the recognition, binding and orientation of RNA in ApaH and their suggested counterparts in ALPH1. **B)** Top, the nucleoside binding site (box) of ApaH accepts RNA in two different orientations depending on the N0 and N1 identity, with preference for A over G, regardless of the downstream nucleotides. Bottom, the possible orientation of substrates in the nucleoside binding site of ALPH1 as inferred from the cleavage patterns in our HPLC assay. For dinucleotides, the nucleoside binding site is preferably occupied by A or m^7^G, and only in a small fraction by G. For longer oligonucleotides and RNAs, the substrates are all bound with N1 in the nucleoside binding site of ALPH1, in agreement with the preferences of the bacterial enzyme.

### Unfit for purpose or just missing partners?

The finding that ALPH1 prefers RNAs (and cap analogues) without the m^7^G methylation is unexpected. One explanation would be, that this substrate preference serves to remove incompletely processed mRNAs, as a quality control. However, in trypanosomes, the enzymatic activity that adds the cap (RNA guanylyltransferase) and the one that adds the m^7^G methylation (guanine-N7-methyltransferase) reside on the same enzyme (45–47), and it is thus unlikely that mRNAs without the m^7^G methylation are frequent by-products of capping. An alternative explanation would be, that an m^7^G-demethylation reaction precedes the decapping, and thus primes the mRNA substrate more sensitive to ALPH1. We cannot exclude such a scenario, but we do consider it unlikely, since we do not find any potential m^7^G-demethylase among ALPH1 interactors (11) and, to the best of our knowledge, no such demethylase has been described in other systems. A third explanation is, that the recruitment of ALPH1 for mRNA decapping may have been a relatively recent invention in trypanosomes and, ALPH1 simply lacked the time to finetune its substrate preferences, also considering that erroneous cleavage of unmethylated RNA caps may not pose a significant problem. Consistently, phylogenetic reconstruction suggests that the loss of DCP2 and gain of ALPH1 is restricted to the subclass of Metakinetoplastina, that encompasses trypanosomatids and bodonids. In the second subclass Prokinetoplastina, represented by Perkinsela and Papus, as well as in euglenids and diplonemids, DCP2-like proteins can be detected by blast searches, while ALPH1 homologs are absent. The promiscuity if the bacterial ApaH towards all Np4Ns and Np4N-capped RNA also suggests that this enzyme family is originally robust in its activity and selectivity.

A fourth explanation, not mutually exclusive with the others, would be that ALPH1 requires activation by other components of the decapping complex, for efficient binding of capped substrates and selective deployment of decapping activity. Decapping is a powerful and terminal determinant in mRNA decay which necessitates tight control to prevent unspecific mRNA degradation. Notably, several mechanistic layers for harnessing DCP2 decapping activity have been uncovered, involving interactions with several activating proteins, as well as autoinhibition (48). The same context renders the low decapping activity of ALPH1 catalytic domain less surprising. In fact, the isolated catalytic domain of yeast DCP2 also displays low basal decapping activity, which is enhanced by the N-terminal, regulatory domain (RD) by two orders of magnitude (49–51) and further by a complex network of decapping activators, including Dcp1, Edc1, Edc2, Edc3 (26, 52) and in mammals Edc4. Specifically, the RD complements the cap binding site and this assembly is stabilized by stepwise recruitment of DCP1 and Edc1 (53). Notably, this process effects DCP2 substrate selectivity by favouring capped RNAs over 5’-triphosphate uncapped RNAs that are preferentially turned over by the isolated catalytic domain (54). Edc3 binding to the CD plays a role in enhancing RNA binding (53) and human EDC4 was suggested to serve as a scaffold, consolidating the DCP1/2 interaction and offering a Xrn1 binding site (55). ALPH1 also assembles into a decapping complex (11). Apart from the Xrn1 homolog XrnA, the ALPH1 core interactors appear to be kinetoplastida specific and remain functionally uncharacterized. It is very likely that these interactors play similar roles in harnessing and modulating of decapping activity.

## Conclusion

We here show that ALPH1 has unique and unexpected preferences for its substrates and cleavage sites, suggesting that the enzyme is rather poorly adapted to its function in decapping trypanosome mRNAs. Whether this poor adaptation is sufficient for its function, or whether regulation by ALPH1 interaction partners within the decapping complex modifies these features, remains a major topic for future studies. The highly unusual features of ALPH1 render the enzyme also an interesting candidate for numerous biotechnology applications, ranging from the production of diphosphate RNA to the possibility to distinguish m^7^G methylated from non-methylated caps by distinct decapping kinetics.

## MATERIAL AND METHODS

### Recombinant ALPH1 protein variants

All ALPH1 variants (ALPH1ΔN, ALPH1ΔNΔC) were produced in *E. coli*, with either 6xHis or GST at the N-terminus for purification. For Figure 2, 4, S1A and S2 ALPH1ΔN was expressed and purified as previously described (4); briefly the proteins were expressed with a tag consisting of a 6xHisTag, a Thrombin cleavage site, a SUMO tag and a TEV cleavage site, purified using standard procedures and the buffer was exchanged to PBS with a PD10 desalting column (Sephadex^TM^ G25M, GE Healthcare); the tag was not removed. For Figure 3, 5BC and 6AB, the His-ALPH1ΔN was purified under phosphate-free conditions from Rosetta (DE3) competent cells. Briefly, the bacterial cell pellet was resuspended in lysis buffer (20 mM Hepes pH8.0, 50 mM NaCl, 5 mM imidazole, 10% glycerin) and cells disrupted by lysozyme (1 mg/ml, 30 min on ice) followed by sonication (6 x 10 sec) and centrifugation (30 min, 4°C, 12000g). The supernatant was incubated with Ni-NTA agarose for 60 minutes and washed twice with wash buffer (20 mM Tris, 300 mM NaCl) containing 5 mM imidazole on a column. The protein was eluted with wash buffer containing 250 mM imidazole and rebuffered to wash buffer using PD10 desalting columns (Sephadex^TM^ G25M, GE Healthcare); the tag was not removed.

ALPH1 variants used for m^7^GTP-pyrene activity assays were produced in *E.coli* with N-terminal 6xHis-SUMO tag. Briefly, the bacteria pellet was resuspended in HisTrap A buffer (50 mM Tris pH 7.5, 500 mM NaCl, 10% glycerin, 20 mM imidazole, 0.5 mM Tris(2-carboxyethyl)phosphin-hydrochlorid (TCEP) and lysed using EmulsiFlex-C3 homogenizer (Avestin), followed by centrifugation (20 min, 4°C, 40000g). The supernatant was applied on a HisTrap HP 5 ml column using AKTA Pure system equilibrated in HisTrap A Buffer. The column was washed with 7CV of HisTrap A buffer and the protein was eluted with a 20-620 mm imidazole gradient in the same buffer. The elution fractions were pooled, diluted 5 times with dilution buffer (50 mM Tris, 10% glycerin, 0.5mM TCEP) and applied on the HiTrap Q HP 5 mL column (ALPH1ΔN) or HiTrap Heparin HP 5 ml column (ALPH1ΔNΔC) equilibrated in HiTrap buffer A (50 mM Tris, 100 mM NaCl, 10% glycerin, 0.5 mM TCEP). The column was washed with 7CV of HiTrap buffer A and the protein was eluted using a 0-100% gradient of HiTrap buffer B (Buffer A supplemented with 500 mM NaCl). The elutions were pooled again, supplemented with 75 ug/mL of SUMO protease (lab-made) and incubated overnight at 8°C. After cleavage, the protein was concentrated on a 10 kDa MWCO Amicon (Milipore) and injected onto Superdex 200 pg 10/300GL Increase column (Cytiva). The column was washed with 1 CV of SEC Buffer (50 mM HEPES pH 7.5, 5% glycerin, 150 mM NaCl, 0.5 mM TCEP). The elution fractions were pooled and flash-frozen in liquid nitrogen, a Coomassie-stained gel loaded with the different fractions of the purification is shown in (Figure S7 in supplementary material).

### Decapping buffer

Decapping buffer is 50 mM Tris-HCl (pH 7.9), 100 mM NaCl, 10 mM MgCl_2_. In early experiments, 1 mM dithiothreitol (DTT) was present, but we found no difference and ommited it in later experiments. For the HPLC assays, the pH was 7.5. Assays from Figure 7D were done in 50 mM Tris-HCl pH 7.5; 50 mM NaCl; 0.5 mM TCEP instead of in decapping buffer.

### RNA (oligos)

Capped RNA oligos were bought from Bio-Synthesis, Texas, USA. To produce the 150 nt long RNAs with and without the m^7^ methylation used in Figure 5A, RNAs were produced by in vitro transcription (IVT) using the HiScribe T7 High Yield RNA Synthesis kit (New England Biolabs) according to the manufacturer instructions. The template for the IVT was a 171 nts long PCR product with pCMV-CLuc 2 as a template and the oligos 5′-ATTAATACGACTCACTATAGG-3′ and 5′-TTAGCTTCACAGGAAGTTG-3′. IVT-produced RNAs were purified with the Monarch Spin RNA Cleanup Kit (New England Biolabs) and subsequently capped with the vaccinia capping system (New England Biolabs), according to the manufacturer’s instructions, in the presence (m7G cap) and absence (G cap) of S-adenosylmethionine (SAM). The capped RNAs were purified using the Monarch Spin RNA Cleanup Kit (New England Biolabs).

### BAAE assays

Each decapping reaction was done in 10 µl volume with 0.025 µM recombinantly produced ALPH1 enzyme (WT, ΔN or ΔNΔC), 0.39 µM capped RNA oligo, 40 units Ribolock RNAse inhibitor (ThermoFisher Scientific) in decapping buffer. Variations of the decapping buffer were used in some experiments and are indicated in the text. The sample was incubated at 37°C (or the indicated temperatures), usually over a time-course. Control reactions were done without enzymes or with the catalytically inactive mutant (Alph1ΔN*; D278:N). The reaction was stopped by ethanol precipitation: 1 µl 3 M RNAse-free sodium acetate (pH 5.5) (Ambion), 1 µl RNAse-free glycogen (ThermoFisher Scientific) and 30 µl cold 100% ethanol was added, the sample incubated at −20°C for 30 minutes, followed by centrifugation (15 minutes, 20,000 g). The pellet (containing the RNA oligo) was washed once in 80% ethanol, air-dried, resolved in 5 µl RNAse-free water (ThermoFisher Scientific). RNA Samples were prepared for gel loading, separated on urea acrylamide gels that contained acryloylaminophenyl boronic acid and imaged on the iBright^TM^ (Invitrogen), as described previously (4). RNA oligos either have a cap 0 or cap 4* (Figure 4A) and usually the sequence of the miniexon. Substrate variations are described in the text.

### Phosphate assays

Phosphate assays were done with ALPH1 enzyme purified without any phosphate. The assay was performed on transparent 96 well plates at 37°C under constant agitation. All substrates were diluted in 1x decapping buffer in 2x concentrations and 25 µl were added to the wells. The reaction was started with 25 µl of a mixture of phosphate-free ALPH1 (variant) in 2 x concentrations and alkaline phosphatase in excess, diluted in decapping buffer. The reaction was stopped at the respective time with 100 µl Biomol®Green (Enzo Life Sciences) and the absorbance was measured at 620 nm in a Tecan Plate reader after 20 minutes. The background (reaction without enzyme) was subtracted. The amount of phosphate was quantified from a phosphate calibration curve. For each experiment, reaction conditions had to be optimized, to be in the (relatively small) linear range of the assay.

### ADP assays

For ADP assays, the decapping reactions were performed in 10 µl volume (in a droplet in a corner of a well from a white low-binding 96 well plate (ThermoFisher, Nunc), at 37° C under constant agitation. The 10 µl contained 5 µl of substrates (at 2x concentrations) and 5 µl of ALPH1 variants (at 2x concentrations), both diluted in 1 x decapping buffer. The reaction was stopped at the required time by adding a mix of 10 μl ADP-Glo™ Reagent and 20 μl Kinase Detection Reagent from the ADP-Glo™ Kinase Assay Kit (Promega). After 30 minutes incubation, luminescence was measured with the Tecan plate reader. The amount of ADP was determined using an ADP calibration curve, done in parallel. Background luminescence was measured from reactions without ALPH1, and subtracted.

### GDP assays

For GDP assays, the decapping reactions were performed in 25 µl volume in white 96 well plates with low-binding (ThermoFisher, Nunc), at 37° C, under constant agitation. The reaction contained 12.5 µl of substrates at 2x concentrations, and 12.5 µl of ALPH1 variant at 2x concentrations, both prepared in 1x decapping buffer. The reaction was stopped by adding 25 µl GDP detection reagent of the GDP-Glo Glycosyltransferase Assay (Promega), prepared according to the manufacturer’s instructions, followed by 30 minutes incubation at room temperature. Luminescence was measured with a Tecan plate reader and the amount of GDP was determined from a GDP calibration curve, recorded in parallel. Background luminescence, measured from reactions without ALPH1, was subtracted.

### Phosphatase assays with pNPP

Phosphatase assays were performed on transparent 96-well plates (ThermoFisher, Nunc). For phosphatase assays with PP1 (Figure 3), the enzyme (Protein Phosphatase-1 Catalytic Subunit (PP1; Sigma-Aldrich)) was incubated for 10 minutes at 30°C in the presence or absence of the respected inhibitors, EDTA (0.3125 mM to 80 mM concentration range), sodium orthovanadate (Na_2_VO_3_; 0.0156 mM to 4 mM concentration range) and okadaic acid (0.3125 µM to 80 µM concentration range) in 1x decapping buffer, in a total volume of 25 µl. Then, 25 µl of 20 mM substrate (pNPP solution in 1x decapping buffer) was added and the absorbance instantly recorded at 405 nm for four minutes, in minute-intervals, at 30° C. The increase in absorbance per minute was calculated and expressed as percentage compared toa non-inhibited control.

For phosphatase assays with ALPH1 variants and alkaline phosphatase (Figure S3), ALPH1 variants (with and without Mn^2+^) and fast alkaline phosphatase (Thermo Scientific) were diluted to 2x concentrations in 1x decapping buffer and added to the microplate in 25 µl volumes. The reaction was started by the addition of 25 µl substrate solution (20 mM of pNPP) in 1 x decapping buffer and the increase in absorbance was immediately measured at 405 nm, at 37°C, for 10 minutes in minute intervals.

### m^7^GTPC_4_-Pyrene assay

For the inhibitor studies (Figure 3B) 50 µl of ALPH1ΔN solution (1 µM) in 1x decapping buffer was incubated for 10 min at 37°C, with 25 µl of inhibitor solution in 1x decapping buffer in black 96-well plates (Thermo Scientific). Then, 25 µl substrate (m^7^GTPC_4_-Pyrene, 800 nM) (14) was added and the fluorescence measured with a Tecan Plate reader, with excitation at 345 nm and emission at 380 nm, for 10 minutes, in two minute intervals. The increase in fluorescence per minute was calculated and compared to a non-inhibited control.

For the assays in Figure 7D, 100 ul assay reactions were prepared as follows: 5 µM m^7^GTP-pyrine probe was added to assay buffer (50 mM Tris-HCl pH 7.5; 50 mM NaCl; 0.5 mM TCEP; 1 mM MgCl_2_) and incubated at 37°C for 10 min. ALPH1 protein was added at the respective concentrations and the reaction was mixed. Fluorescence signal was detected in 1 min intervals, with a TECAN plate reader (λexc=345nm, λem= 380nm) at 37°C. Each reaction signal was normalized to the negative control (without ALPH1 protein) and the linear range of the reaction was determined. Within this range, the initial reaction rate was calculated with the linear regression method.

### HPLC assay

The reaction mixture (100 μL) consisting of cap analog (50 μM), ALPH1ΔN (1 μM) and buffer (50 mM TRIS·HCl pH 7.5, 100 mM NaCl, 10 mM MgCl_2_) was incubated at 25°C on a shaker (250 rpm) for 1 h. Then, a suspension of Chelex® 100 sodium form (50-100 mesh) in 50 mM AcONH_4_ pH 5.9 (washed several times to get pH∼8) was added and the sample was shaken gently for 20 min. The mixture was centrifuged in Amicon® Ultra Centrifugal Filter, 10 kDa MWCO (14,100 rcf, 5 min), 100 μL water was added to the filter and the mixture was centrifuged again. Combined filtrates (flow-through) were analyzed by RP HPLC (Shimadzu LC-40 XS) on Gemini NX-C18 column (150 x 4.6 mm, 3 μm, 110Å) using a linear gradient elution with 0–15% acetonitrile in 0.05 M ammonium acetate buffer pH 5.9 and the collected peaks were additionally analyzed by mass spectrometry (AB Sciex API 3200).

### Synthesis of m^7^GpppA_m_UUUU and GpppA_m_UUUU

#### Solid-phase synthesis of pA_m_UUUU

Oligonucleotide 5′-phosphate pA_m_UUUU was synthesized by the phosphoramidite method on solid phase (Primer Support 5G, ribo U 300, Cytiva; 50 μmol scale) using an ÄKTA oligopilot plus synthesizer. In the coupling step, 5 equivalents of 3′-O-phosphoramidate [2′-OMe-A^Pac^, 2′-OTBDMS-U or Bis(2-cyanoethyl)-N,N-diisopropyl phosphoramidite] and 0.30 M 5-(benzylthio)-1-*H*-tetrazole in acetonitrile were recirculated through the column for 15 minutes. A solution of 3% (v/v) dichloroacetic acid in toluene was used as a detritilation reagent and 0.05 M iodine in pyridine for oxidation, 20% (v/v) *N*-methylimidazole in acetonitrile as Cap A and a mixture of 40% (v/v) acetic anhydride and 40% (v/v) pyridine in acetonitrile as Cap B. After the last cycle of synthesis, RNAs, still on the solid support, were treated with 20% (v/v) diethylamine in acetonitrile. The product was released from the solid support and base protecting groups were removed using 5 mL of AMA (33% aqueous ammonia + 40% aqueous methylamine 1:1; 30°C, 2h) and TBDMS groups were removed by incubation of oligo dissolved in DMSO (200 μL) with triethylammonium trihydrofluoride (250 μL) and triethylamine (430 μL) at 65°C for 2.5 h. The product was isolated as a triethylammonium salt (1910 mOD_260nm_, 42.9 μmol) by ion-exchange chromatography on DEAE Sephadex (gradient elution with 0–1.2M triethylammonium bicarbonate).

#### Activation of 5′-phosphate (synthesis of Im-pA_m_UUUU)

Triethylammonium salt of pA_m_UUUU (1910 mOD_260nm_, 42.9 µmol) was dissolved in anhydrous dimethylformamid (DMF;860 µL) followed by the addition of imidazole (46.7 mg, 686 µmol), triethylamine (36 µl, 257 µmol), 2,2’-dithiodipyridine (56.6 mg, 257 µmol) and triphenylphospine (67.4 mg, 257 µmol). After 24 h, the product was precipitated with a cold solution of NaClO_4_ (42.0 mg, 343 µmol) in acetonitrile (15 mL). The precipitate was centrifuged (5000 rpm, 8 min) in a 50 ml conical tube at 4°C, washed with cold acetonitrile by centrifugation 3 times, and dried under reduced pressure to give Im-pA_m_UUUU (62 mg, 35.5 μmol, 83%) as a white solid.

#### Synthesis of GpppA_m_UUUU

Im-pA_m_UUUU (5.0 mg, 2.9 μmol) and triethylammonium salt of guanosine 5′-diphosphate (GDP, 4.6 mg, 7.2 µmol) were suspended in anhydrous DMF (115 µL) and anhydrous ZnCl_2_ (3.9 mg, 29 µmol) was added. The mixture was stirred at room temperature for ca. 2.5 h and the reaction was quenched by addition of 0.9 mL of an aqueous solution of EDTA (20 mg/mL) and NaHCO_3_ (10 mg/mL). The product was isolated by semi-preparative RP HPLC (gradient elution 0–15% acetonitrile in 0.05 M ammonium acetate buffer pH 5.9) to afford – after evaporation and repeated freeze-drying from water – an ammonium salt of GpppA_m_UUUU (78 mOD, 1.4 µmol, 50%) as a white solid.

#### Synthesis of m^7^GpppA_m_UUUU

Im-pA_m_UUUU (6.6 mg, 3.8 μmol) and triethylammonium salt of 7-methylguanosine 5′-diphosphate (m^7^GDP, 5.0 mg, 7.6 µmol) were suspended in anhydrous DMF (1.5 mL) and anhydrous ZnCl_2_ (4.1 mg, 30 µmol) was added. The mixture was stirred at room temperature overnight and the reaction was quenched by addition of 0.6 mL of an aqueous solution of EDTA (20 mg/mL) and NaHCO_3_ (10 mg/mL). The product was isolated by ion-exchange chromatography on DEAE Sephadex using a linear gradient of TEAB (0–1.2M) and purified by semi-preparative RP HPLC (gradient elution 0–15% acetonitrile in 0.05 M ammonium acetate buffer pH 5.9) to afford – after evaporation and repeated freeze-drying from water – ammonium salt of m^7^GpppA_m_UUUU (6.6 mg, 3.1 µmol, 82%) as a white solid.

### Trypanosome cell culture and transgenic cell lines

Trypanosoma cells Trypanosoma brucei Lister 427 procyclic cells in logarithmic growth were used for all experiments. Cells were grown in SDM-79 supplemented with 5% (v/v) FCS and 75 μg/ml hemin at 27°C, 5% CO2, and appropriate drugs. Transgenic trypanosomes were generated by standard procedures (56).

For the auxin inducible degron system of ALPH1, the auxin-mother cell line (which constitutively expresses OsSKP1, OsCUL1, OsRBX1, OsTIR1(F74G) and OsARF (22) was modified: one allele of ALPH1 was replaced by a phleomycine resistance marker and the second allele was replaced by a C-terminal fusion of ALPH1 to OsAID2-3HA (puromycin selection), both via pPOT system for genetic manipulation via a PCR transfection (57). For the rescue experiments, ALPH1 variants (wt, wt* (inactive),ΔN, ΔC, ΔNΔC) were integrated to the tubulin locus for constitutive expression. Protein degradation was induced by adding 50 µM 5-Ph-IAA (MedChem Express, HY-134653) to the cells, from a 50 mM stock (in DMSO, stored at −80°C).

### Western blot

The western blot was performed using standard procedures and probed with anti-ALPH1 (polyclonal, anti-rabbit, 1:1000) and anti-HA (monoclonal anti rat, 3F10, Roche, 1:1000). IRDye680CW goat anti-rabbit and IRDye 800CW Goat anti-Rat IgG (Licor, 1:20,000) were used as secondary antibodies.

### 3D model preparation and visualisation

PDB entry 9ooy was used for the model of *E.coli* ApaH with GppppAU bound. The predicted model of ALPH1 was obtained using Alphafold 3 webserver (58) accessed on 21.01.2026. Structural models were aligned and visualised using ChimeraX v.1.10 (59).

## SUPPLEMENTARY DATA

Figure S1: control experiments to mRNA decapping assays (to Figure 1)

Figure S2: control experiments for basic enzyme characterisation (to Figure 2)

Figure S3: phosphatase activity of ALPH1ΔN

Figure S4: BAAE assays showing decapping activity of ALPH1ΔNΔC

Figure S5: substrate preferences of ALPH1ΔNΔC

Figure S6: replicate for Figure 5D

Figure S7: Coomassie-stained gels showing the purification of the ALPH1 variants used for the m^7^GTP-pyrene activity assays.

## ACKNOWLEDGEMENT

This worked was supported by a trilateral DFG/GACR/NCN (KR4017/12-1; 24-14298L; National Science Centre, Poland grant 2022/04/Y/NZ1/00114 WEAVE-UNISONO) grant to SK, MZ and MG. MW received financial support from the National Science Centre, Poland (2022/47/D/ST4/00386). In addition, SK was funded by the DFG grants KR4017/4-1 and KR4017/4-2 and SK and FH were supported by a bilateral DAAD and PROBRAL/CAPES grant [DAAD Projekt-ID: 57597990; PROBRAL/CAPES 88881.628073.2021-01]. Markus Krischke and Martin Müller, both University of Würzburg, Germany, are thanked for their help with initial HPLC experiments. Carolin Schlitz, University of Würzburg, is acknowledged for help with Euglena phylogenetic studies.

