## Supplementary Figures for "The trypanosome mRNA decapping enzyme ALPH1 prefers caps without m^7^G methylation and produces diphosphate-RNA"

### Figure S1

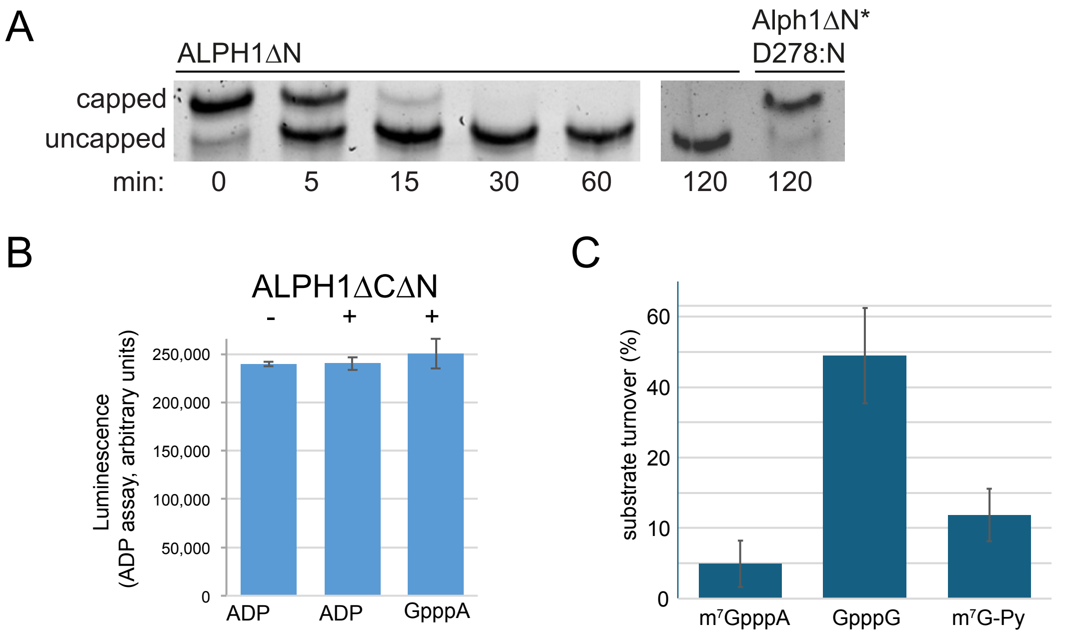


**Figure S1: Control experiments to Figure 1 (mRNA decapping assays)**

**A) An inactive ALPH1 mutant does not cause a band-shift in BAAE**
An 39 nucleotide long RNA oligo with the miniexon sequence and a cap 0 was incubated with recombinant ALPH1ΔN in decapping buffer over a time-course. Capped and uncapped oligo are separated by boronate affinity acrylamide electrophoresis. No decapping is observed at incubation with Alph1ΔN* (an inactive mutant, D278:N).
**B) ADP is not a substrate for ALPH1**
125 µM ADP was incubated without and with ALPH1ΔCΔN. ADP was measured with the ADP assay and is shown in arbitrary units. The addition of ALPH1ΔN does not reduce the amount of ADP, proving that ADP is not a substrate for ALPH1. As a positive control for ALPH1 activity, 125 µM GpppA was used, which is completely cleaved to ADP (detected) and GMP (not detected in this assay).
**C) $m^7$GTPC4-Pyrene is accepted as a substrate by ALPH1**
mRNA decapping was measured using the phosphate assay. 1.5 nM His-tagged recombinant ALPH1ΔN was incubated with 50 µM of substrates for 25 minutes at 37°C. Substrate turnover is presented, with error bars representing standard deviations from three replicate experiments.

A

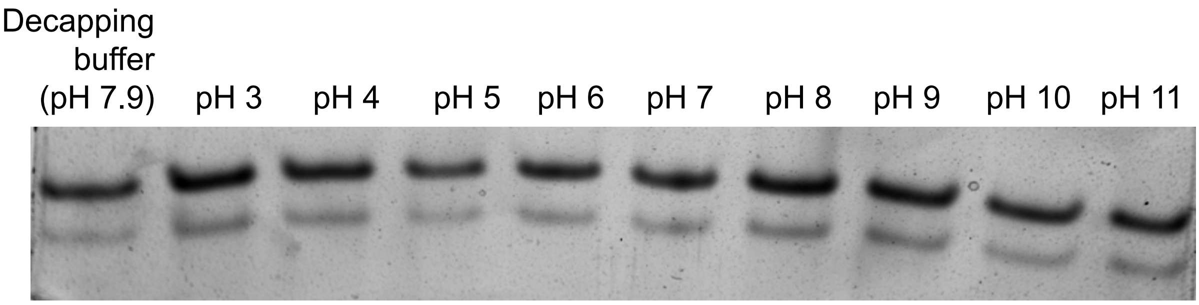


B

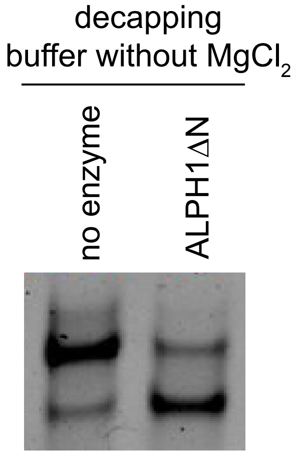


**Supplementary Figure S2**

**A)** A BAAE decapping assay was performed in decapping buffer at different pH values, without the addition of ALPH1 enzyme, using m7GpppA-ME (cap 0) as a substrate. No decapping was observed in the absence of ALPH1, proving that changes in pH values does not cause decapping.

**B)** A BAAE decapping assay was performed with ALPH1ΔN in decapping buffer without magnesium ions. Data of one representative experiment out of > five experiments is shown.

### Figure S3

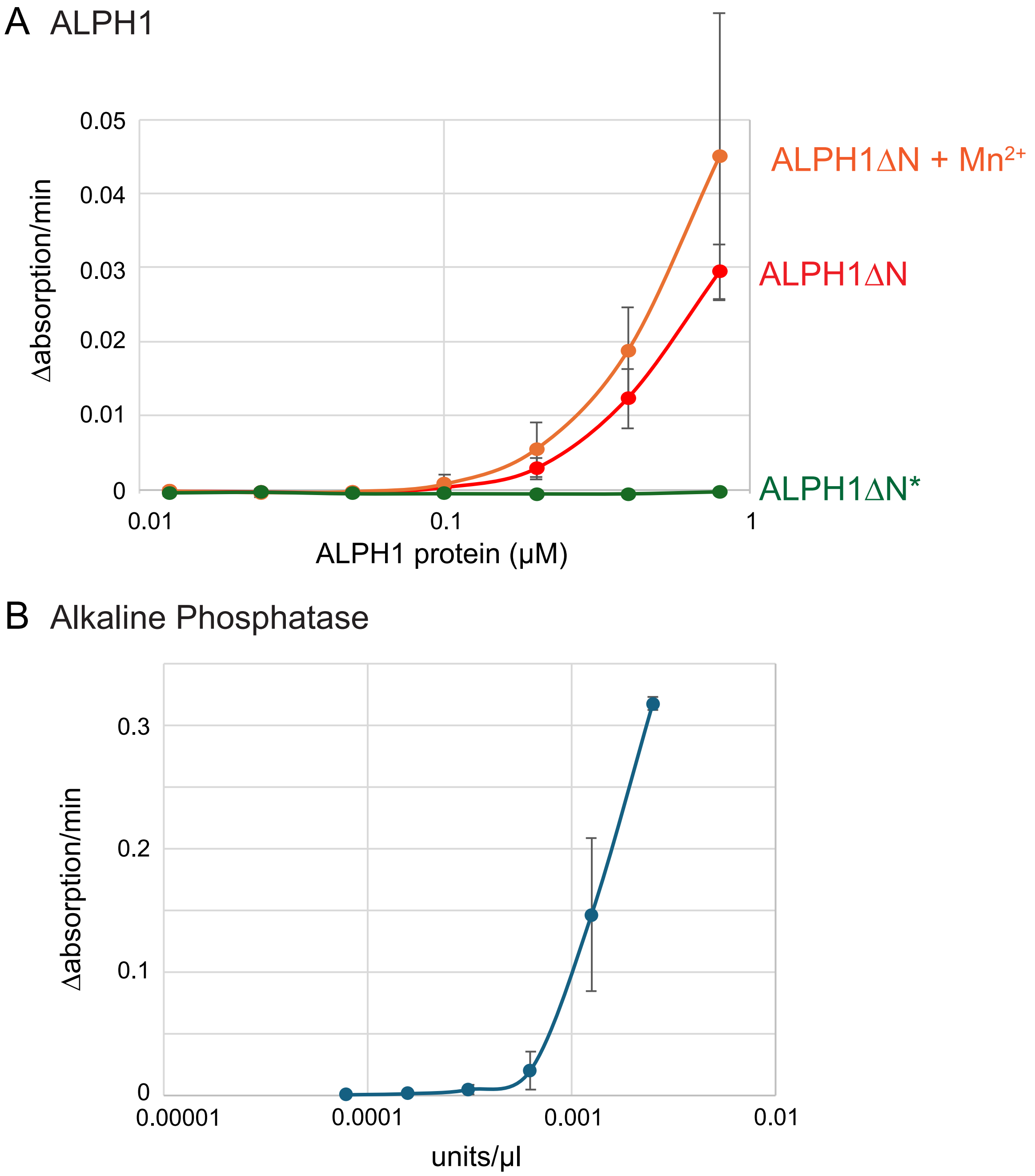


**Figure S3: Phosphatase activity of ALPH1**

**A)** pNPP turnover by up to 0.8 μM ALPH1ΔN was measured over 10 minutes in decapping buffer, with and without manganese (1 mM). The catalytically inactive mutant ALPH1ΔN* ((D278:N) served as a negative control.

**B)** As a positive control, the activity of Alkaline Phosphatase was measured.

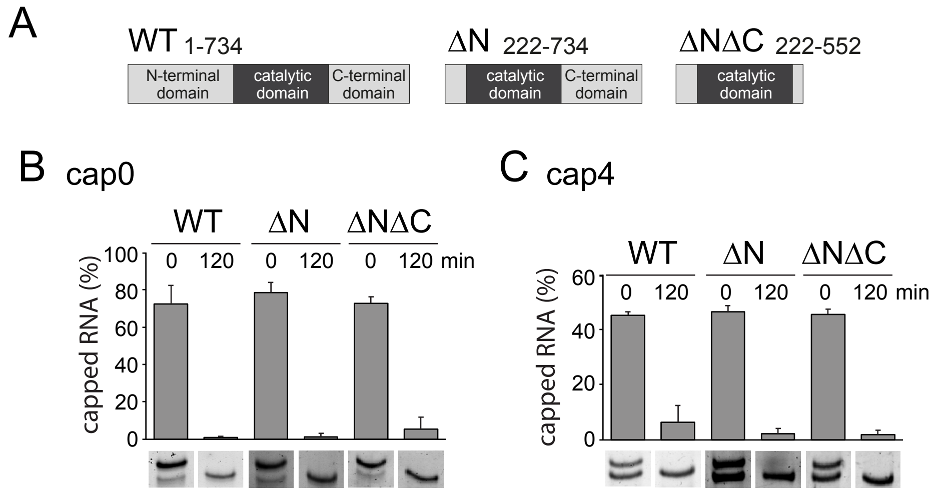


**Figure S4: The catalytic domain of ALPH1 has mRNA decapping activity**

**A)** To scale schematics of the ALPH1 truncations used in this experiment.

**B)** A BAAE mRNA decapping assay was done for120 minutes using the three ALPH1 variants shown in A. The substrate was the 39 nucleotide ME RNA capped with either cap0 or cap4* (compare Figure 4). The percentage of capped RNA was quantified from three independnet experiments and the average values are shown as bars with error bars representing the standard deviation.

### Figure S5

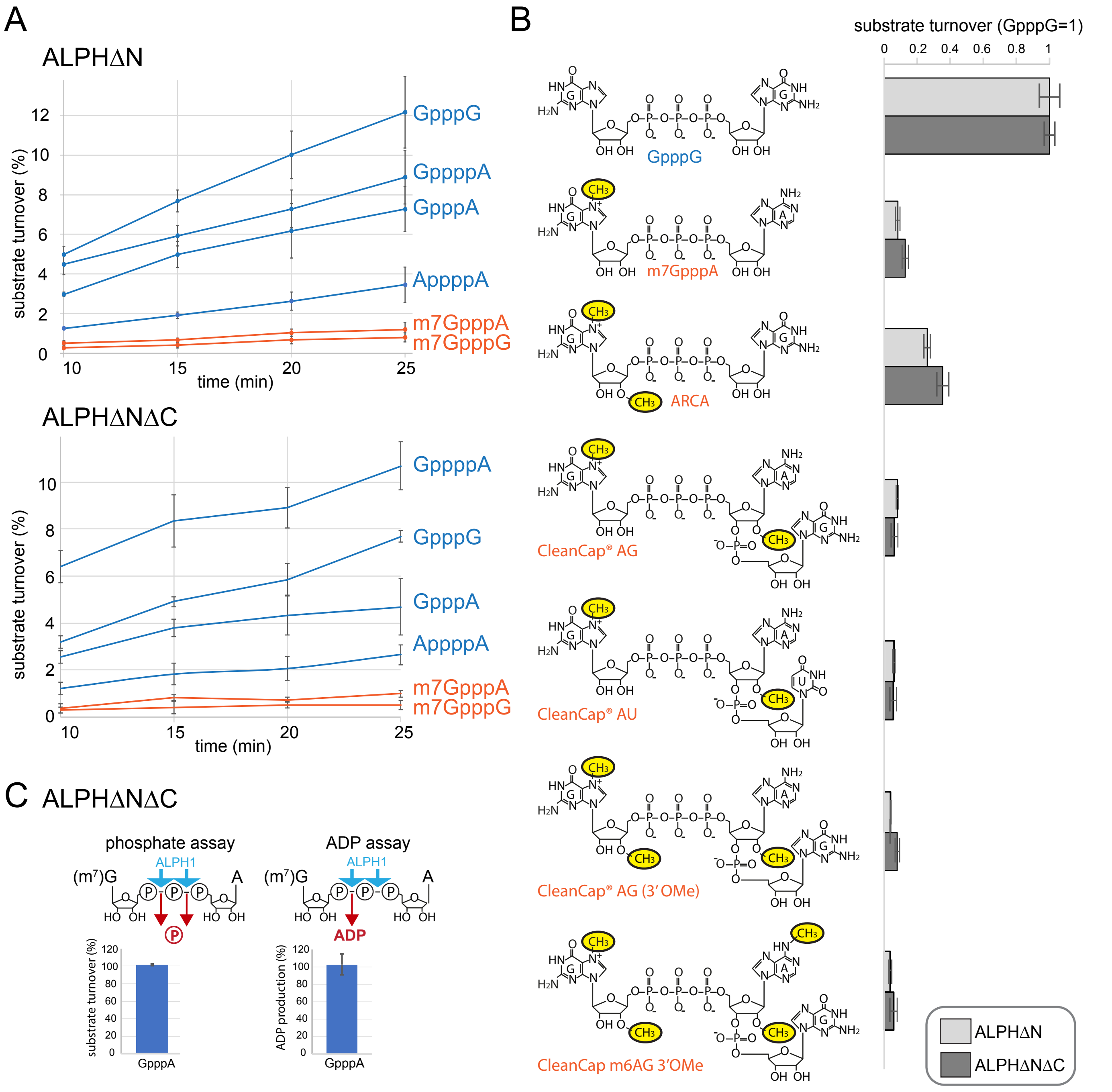


**Figure S5: ALPH1ΔNΔC has similar substrate preferences to ALPH1ΔN**

**A and B)** The experiment shown in Figure 5 was repeated with ALPH1ΔNΔC. The conditions are identical, except that ALPH1ΔNΔC was used at 15 nM, thus, at a ten fold higher concentration than ALPH1ΔN, to compensate for its lower activity. The ALPH1ΔN data of Figure 5 are included for comparison.

**C)** The cleavage site of GpppA was determined by combining the phosphate assay (to ensure completion) with the ADP assay, exactly like in Figure 6A. The assay was done with 3.9 μM substrate and 0.12 μM ALPH1ΔNΔC for 60 minutes.

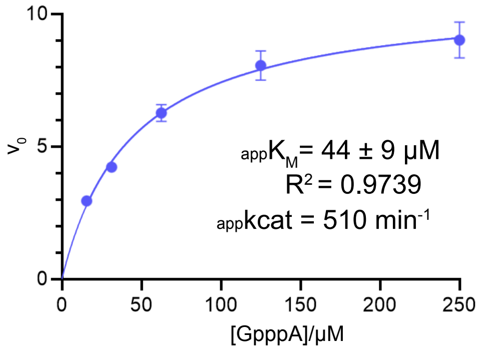


**Figure S6: Replicate of Figure 5D**

Determination of the apparent $K_M$ ($_{app}K_M$) for ALPH1ΔN (21 nM) for the GpppA substrate via phosphate assay. The multiple-turnover reaction was over 15 min and $_{app}K_M$, $_{app}kcat$ and $R^2$ values are indicated below the curves.

Figure S7

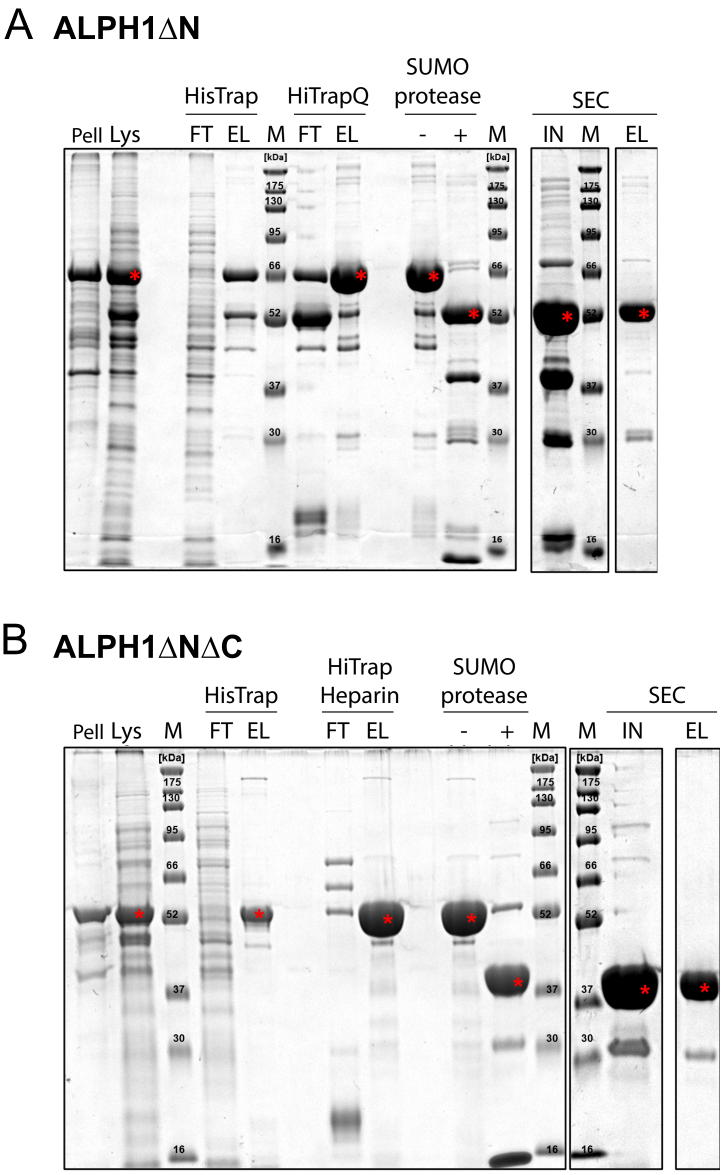


**Figure S7: Coomassie-stained gels to control the purification of ALPH1 variants**

SDS gels were loaded with samples taken during the purification of ALPH1ΔN (**A**) and ALPH1ΔNΔC (**B**) and proteins were detected by Coomassie staining.

Pell=pellet; Lys=lysate; FT=flow through; EL=eluate; M=marker; IN=Input. The red star marks the band referring to the ALPH1 variant. For further details, refer to the material and method section.
